# Unique features of stimulus-based probabilistic reversal learning

**DOI:** 10.1101/2020.09.24.310771

**Authors:** Carl Harris, Claudia Aguirre, Saisriya Kolli, Kanak Das, Alicia Izquierdo, Alireza Soltani

**Affiliations:** Department of Psychological and Brain Science, Dartmouth College, Hanover, New Hampshire, United States 03755; Department of Psychology, University of California-Los Angeles, Los Angeles, California, United States 90095; The Brain Research Institute, University of California-Los Angeles, Los Angeles, California, United States 90095; Integrative Center for Learning and Memory, University of California-Los Angeles, Los Angeles, California, United States 90095; Integrative Center for Addictions, University of California-Los Angeles, Los Angeles, California, United States 90095

**Keywords:** reversal learning, probabilistic learning, win-stay, lose-shift, stimulus-reward, reinforcement learning, orbitofrontal cortex, DREADDs

## Abstract

Reversal learning paradigms are widely-used assays of behavioral flexibility with their probabilistic versions being more amenable to studying integration of reward outcomes over time. Prior research suggests differences between initial and reversal learning, including higher learning rates, a greater need for inhibitory control, and more perseveration after reversals. However, it is not well-understood what aspects of stimulus-based reversal learning are unique to reversals, and whether and how observed differences depend on reward probability. Here, we used a visual probabilistic discrimination and reversal learning paradigm where male and female rats selected between a pair of stimuli associated with different reward probabilities. We compared accuracy, rewards collected, omissions, latencies, win-stay/lose-shift strategies, and indices of perseveration across two different reward probability schedules. We found that discrimination and reversal learning are behaviorally more unique than similar: latencies to select the better option, win-stay strategies, and perseveration were of a different pattern in reversal learning compared to discrimination learning. Additionally, fit of choice behavior using reinforcement learning models revealed a lower sensitivity to the difference in subjective reward values (greater exploration) and higher learning rates for the reversal phase. Interestingly, a consistent reward probability group difference emerged with a richer environment associated with longer reward collection latencies than a leaner environment. We also replicated previous reports on sex differences in reversal learning. Future studies should systematically compare the neural correlates of fine-grained behavioral measures to reveal possible dissociations in how the circuitry is recruited in each phase.

## Introduction

A critical feature of goal-directed, instrumental behavior is the ability to discriminate stimuli that predict reward from those that do not, and further, to flexibly update the response to those stimuli if predictions become inaccurate. Discrimination learning paradigms in rodents often involve pairing of an action (e.g., lever pressing, digging in a bowl, nosepoking stimuli on a touchscreen, or displacing an object) with an outcome (e.g., a desirable food reward). In typical paradigms, two or more stimuli are presented concurrently and the subject learns about the features of the stimuli that bring about reward and those that do not (e.g., nosepoking S_A_ results in a better probability of reward than S_B_; pressing left lever yields better payout than right lever; scent A is more rewarded than scent B) (Alvarez & Eichenbaum, 2002; Dalton, Wang, Phillips, & Floresco, 2016; Eichenbaum, Fagan, & Cohen, 1986; Izquierdo et al., 2013; Schoenbaum, Chiba, & Gallagher, 2000; Schoenbaum, Nugent, Saddoris, & Setlow, 2002). With training, subjects become increasingly proficient at discrimination, in line with the associative rules imposed by the experimenter. The stimulus-reward rules can be deterministic (e.g., S_A_ results in a sucrose pellet reward and S_B_ does not) or probabilistic (e.g., S_A_ results in a better probability of reward over S_B_), with deterministic and probabilistic schedules of reinforcement producing marked dissociations in the neural circuity recruited (Averbeck & Costa, 2017; Costa, Dal Monte, Lucas, Murray, & Averbeck, 2016).

In reversal learning paradigms (Izquierdo, Brigman, Radke, Rudebeck, & Holmes, 2017; Izquierdo & Jentsch, 2012), after either reaching a discrimination learning criterion for accuracy (Brushfield, Luu, Callahan, & Gilbert, 2008; Izquierdo et al., 2013; Stolyarova et al., 2019), a number of consecutive correct responses (Dalton, Phillips, & Floresco, 2014; Dalton et al., 2016), or a fixed block length of trials (Farashahi et al., 2017; Soltani & Izquierdo, 2019), the stimulus-reward contingencies are reversed. At reversal, the trained response no longer results in a better probability of reward, though it usually remains the more frequently chosen option because of initial discrimination training. Indeed, usually reversals are acquired more slowly than the original discrimination, and younger subjects are quicker to learn than older ones (Brushfield et al., 2008; Schoenbaum, Setlow, Saddoris, & Gallagher, 2006). Perhaps partly due to this difference with original learning, reversal learning is considered unique in its requirement of flexibility, because it involves the subject inhibiting the prepotent response and, instead, responding to stimuli that were previously irrelevant. Other popular views are that discrimination and reversal learning phases occupy different task “spaces” (Wilson, Takahashi, Schoenbaum, & Niv, 2014) and that the phases differ in the likelihood that changes in contingencies will occur (Jang et al., 2015). Importantly, probabilistic learning and reversal paradigms, in particular, are more amenable to the application of reinforcement learning (RL) models that can estimate parameters for choice behavior based on the integration of previous rewarded and non-rewarded trials (Lee, Seo, & Jung, 2012). Despite various favored accounts, it is not well-understood how *behaviorally unique* reversal learning is compared to original (initial discrimination) learning. For example, latency measures can be used to dissociate attention, decision speed, and motivation via analyses of the time taken to initiate trials, choose a stimulus, and collect reward, respectively (Aguirre et al., 2020). Here, we explored if such detailed trial-by-trial measures of latencies and omissions differ across learning phases. Further, we probed if there are differences between reward probability schedules on these various measures by comparing the ability of separate cohorts of animals tested on different reward probability schedules to discriminate between the better and worse options, which were rewarded with a probability of 0.90 vs. 0.30, compared to 0.70 vs. 0.30. We also analyzed win-stay and lose-shift strategies, and perseveration and repetition metrics in each learning phase. Finally, we investigated if RL models fit these learning phases differently. We studied this in both male and female animals.

We found a higher perseveration index and reduced use of win-stay strategies unique to early reversal compared to discrimination learning, which was expected. However, more surprisingly, we found the most robust, consistent measures across discrimination and reversal learning phases to be latencies to choose the better option and to collect reward. These metrics are proxies for decision speed and motivation, respectively. As for RL models, we found a lower sensitivity to the difference in subjective reward values and higher learning rates during reversal than initial discrimination. There were also sex differences, particularly in reversal learning, which replicate our previous findings using this paradigm. Interestingly the only consistent probability group difference we observed across discrimination and reversal learning phases was in motivation (i.e. a richer environment was associated with longer reward collection latencies than a leaner environment). Collectively, our fine-grained analyses suggest that trial-by-trial behavioral measures of latencies and strategies may be particularly sensitive metrics to pair with neural correlate data in reversal learning. These measures may also be revealing in uncovering the unique substrates of flexible learning.

## Methods

### Subjects

Subjects were N=25 adult male (n=13) and female (n=12) Long-Evans rats (Charles River Laboratories) aged > post-natal-day (PND) 60 at the start of testing. Rats arrived to the vivarium between PND 40-60. The rats included in this report served as controls for two different experiments: one was a cohort of water (H_2_O)-only drinking (n=8 female and n=8 male) rats that served as controls in an ethanol study (see *2-Bottle Choice*) and the other was a cohort of rats (n=5 female and n=4 male) that experienced surgical procedures (see *Surgery*), serving as controls for a study targeting the orbitofrontal cortex (OFC) with DREADDs. Importantly, all rats were the same age (PND 140-155) when pretraining commenced, and further, all rats were part of experiments that ran in parallel, minimizing differences between cohorts.

Before any treatment, all rats underwent a 3-day acclimation period during which they were pair-housed and given food and water *ad libitum*, and remained in cages with no experimenter interference. Following this 3-day acclimation period, animals were handled for 10 min per animal for 5 consecutive days. During the handling period, the animals were also provided food and water *ad libitum*. After the handling period, animals were individually-housed under standard housing conditions (room temperature 22–24° C) with a standard 12 h light/dark cycle (lights on at 6am). Following either 2-bottle choice or surgery, rats were tested on probabilistic discrimination and reversal learning, as below. All procedures were conducted in accordance with the recommendations in the Guide for the Care and Use of Laboratory Animals of the National Institutes of Health and the Chancellor’s Animal Research Committee at the University of California, Los Angeles.

### 2-Bottle Choice

Home cages were modified to allow for the placement of two bottles for drinking, whereas standard housing allows for only one bottle. Rats (n=16; 8 male and 8 female) included in this analysis were singly-housed and given access to 2 H_2_O bottles simultaneously (no ethanol) for 10 weeks, the same duration that experimental animals were provided the choice of ethanol vs. H_2_O. Weight of bottles was measured three times per week to measure consumption amounts, compared to a control cage placed on the same rack to account for leakage. Rats were not monitored for weight during this time.

### Surgery

#### Viral Constructs

Rats (n=9; 4 male and 5 female) included in the present comparison were singly-housed and allowed to express DREADDs or eGFP in OFC for 6 weeks, the same duration experimental animals that were treated with clozapine-N-oxide (CNO) were allowed to express DREADDs. An adeno-associated virus AAV8 driving the hM4Di-mCherry sequence under the CaMKIIa promoter was used to express DREADDs on OFC neurons (AAV8-CaMKIIa-hM4D(Gi)-mCherry, packaged by Addgene) (Addgene, viral prep #50477-AAV8). A virus lacking the hM4Di DREADD gene and only containing the green fluorescent tag eGFP (AAV8-CaMKIIa-EGFP, packaged by Addgene) was also infused into OFC in separate groups of animals as a null virus control. Altogether the animals included in these sets of analyses served as control groups for a larger experiment in which they were given subcutaneous injections of either CNO or a saline vehicle (VEH) prior to reversal learning. The animals included here were 5 animals prepared with hM4Di DREADDs in OFC who received VEH, 2 animals prepared with eGFP in OFC who received VEH, and 2 animals prepared with eGFP in OFC, who received CNO during reversal learning. Importantly, although these animals received virus in OFC, the DREADDs were not activated. Further, we provide analyses to show these subgroups did not differ, and could be combined.

#### Surgical Procedure

Infusion of AAV virus containing DREADD or eGFP (n=9) in OFC was performed using aseptic stereotaxic techniques under isoflurane gas (1-5% in O_2_) anesthesia prior to any behavioral testing experience. Before surgeries were completed, all animals were administered 5mg/kg s.c. carprofen (NADA #141–199, Pfizer, Inc., Drug Labeler Code: 000069) and 1cc saline. After being placed in the stereotaxic apparatus (David Kopf; model 306041), the scalp was incised and retracted. The skull was leveled to ensure that bregma and lambda were in the same horizontal plane. Small burr holes were drilled in the skull above the infusion target. Virus was bilaterally infused at a rate of 0.02 μl per minute for a total volume of 0.2 μl per hemisphere into OFC (AP = +3.7; ML= ±2.0; DV = −4.6, relative to bregma). After each infusion, 10 min elapsed before the syringe was pulled up.

### Food Restriction

Five days prior to any behavioral testing, rats were placed on food restriction with females on average maintained 12-14 grams/ day and males given 16-18 grams/ day of chow. Food restriction level remained unchanged throughout behavioral testing, provided animals completed testing sessions. Water remained freely available in the home cage. Animals were weighed every other day and monitored closely to not fall below 85% of their maximum, free-feeding weight.

### Learning

#### Pretraining

Behavioral testing was conducted in operant conditioning chambers outfitted with an LCD touchscreen opposing the sugar pellet dispenser. All chamber equipment was controlled by customized ABET II TOUCH software.

The pretraining protocol, adapted from established procedures (Stolyarova & Izquierdo, 2017), consisted of a series of phases: Habituation, Initiation Touch to Center Training (ITCT), Immediate Reward Training (IMT), designed to train rats to nosepoke, initiate a trial, and select a stimulus to obtain a reward (i.e. sucrose pellet). During habituation, rats were required to eat five pellets out of the pellet dispenser inside the chambers within 15 min before exposure to any stimuli on the touchscreen. ITCT began with the display of white graphic stimuli on the black background of the touchscreen. During this stage, a trial could be terminated for one of two reasons: if a rat touched the displayed image and received a reward, or if the image display time (40 s) ended, after which the stimulus disappeared, a black background was displayed, and a 10 s inter-trial interval (ITI) ensued. If the rat did not touch within 40 s this was scored as an *initiation omission*. IMT began in the same way as ITCT, but the disappearance of the white graphic stimulus was now paired with the onset of a target image immediately to the left or right of the stimulus (i.e. forced-choice). During this stage, a trial could be terminated for one of three reasons. First, if a rat touched the center display (i.e. white graphic stimulus) and touched the image displayed on either side, after which there was a dispensation of one sucrose pellet and illumination of the tray-light. Second, if the rat failed to touch the center white graphic stimulus after the display time ended (40 s), after which the stimulus disappeared, a black background was displayed, and a 10 s ITI ensued, scored as an *initiation omission*. Third, if the image display time (60 s) ended, after which the stimulus disappeared, a black background was displayed, and a 10 s ITI ensued, scored as a *choice omission*. Rats could also fail to respond to the center stimulus within 40 s during this phase (i.e. initiation omission, as in the previous phase). For habituation pretraining, the criterion for advancement was collection of all 5 sucrose pellets. For ITCT, the criterion to the next stage was set to 60 rewards consumed in 45 min. The criterion for IMT was set to 60 rewards consumed in 45 min across two consecutive days.

#### Probabilistic Discrimination Learning

After completion of all pretraining schedules, rats were advanced to the discrimination (initial) phase of the PRL task, in which they would initiate a trial by touching the white graphic stimulus in the center screen (displayed for 40 s), and choose between two visual stimuli presented on the left and right side of the screen (displayed for 60 s) counterbalanced between trials, assigned as the better or worse options, with a reward (i.e. sucrose pellet) probability of either p_R_(B)= 0.90 or 0.70 (i.e. better option) or p_R_(W)=0.30 (i.e. worse option). If a trial was not initiated within 40 s, it was scored as an initiation omission. If a stimulus was not chosen, it was scored as a choice omission, and a 10 s ITI ensued. If a trial was not rewarded, a 5 s time-out would follow, subsequently followed by a 10 s ITI. Finally, if a trial was rewarded, a 10 s ITI would follow after the reward was collected (**Figure 1**). The criterion was set to 60 or more rewards consumed and selection of the better option in 70% of the trials or higher during a 60 min session across two consecutive days. After reaching the criterion for the discrimination phase, the rats advanced to the reversal phase beginning on the next session. Notably, one animal in the 90-30 group and five animals in the 70-30 group did not meet discrimination criterion and were forced-reversed after 25+ days.

**Figure 1.**
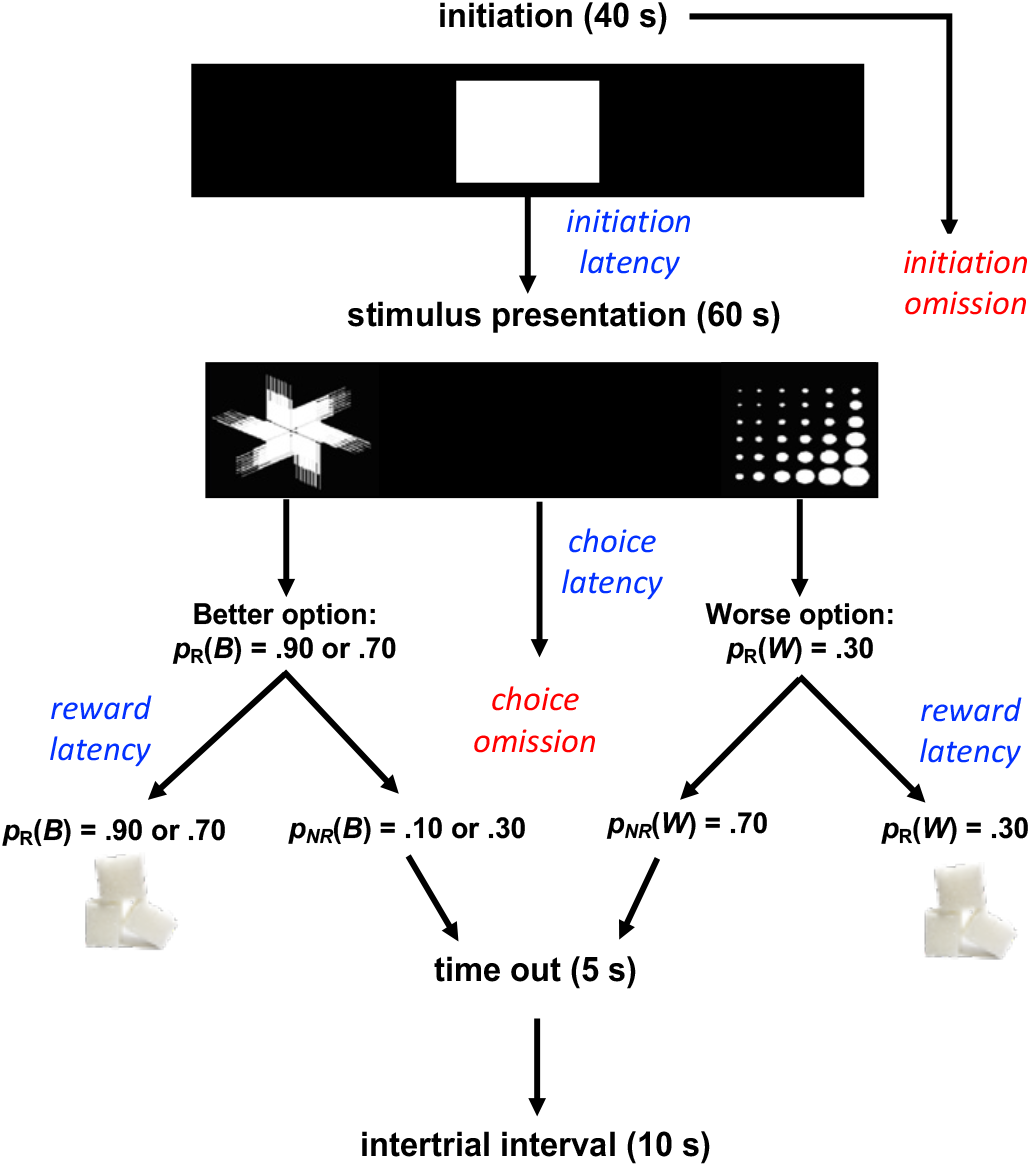
Task design. Schematic of probabilistic learning task. Rats initiated a trial by nosepoking the center stimulus (displayed for 40 s) and then selected between two visual stimuli pseudorandomly that were presented on either the left and right side of the screen (displayed for 60 s), assigned as the better (B) and worse (W) options. Correct nosepokes were rewarded with a sucrose pellet with probability p_R_(B)=0.90 or 0.70 versus p_R_(W)=0.30. If a trial was not rewarded [p_NR_(B) or p_NR_(W)], a 5 s time-out would commence. If a stimulus was not chosen, it was considered a choice omission and a 10 s ITI would commence. Rats could also fail to initiate a trial, in which case, it was scored as an initiation omission. If a trial was rewarded, a 10 s ITI would follow reward collection. Other prominent measures collected on a trial-by-trial basis were trial initiation latency (time to nosepoke the center white square), choice latency (time to select between the two stimuli), and reward latency (time to collect reward in the pellet magazine).

#### Probabilistic Reversal Learning

After the discrimination phase, the rats advanced to the reversal phase during which rats were required to remap stimulus-reward contingencies and adapt to reversals in the reward probabilities. The stimuli associated with the p_R_ (B)=0.90 or 0.70 probability (i.e. better option), would now be associated with a p_R_(W)=0.30 probability of being rewarded (i.e. worse option). Consistent with prior literature showing freely-behaving rodents exhibit slow learning on probabilistic reversals with visual stimuli (Aguirre et al., 2020), most animals from either cohort did not meet a 70% criterion before the termination of the study, so we limited our analyses to the first seven sessions of discrimination and reveral phases for all animals.

### Data Analyses and computational modeling

MATLAB (MathWorks, Natick, Massachusetts; Version R2019b) was used for all statistical analyses; MATLAB and Prism8 were used for figure preparation. Data were analyzed with a series of mixed-effects General Linear Models (GLM); omnibus analyses across discrimination and reversal phases in early learning (operationally defined as the first seven sessions), and then individual analyses within each phase separately if justified by a phase interaction. We and others have analyzed early learning in previous work, as it may be particularly informative to revealing sensitivity to reward feedback and perseveration (Izquierdo et al., 2010; Jones & Mishkin, 1972; Stolyarova, O’Dell, Marshall, & Izquierdo, 2014; Stolyarova et al., 2019), measured using touchscreen response methods (Izquierdo et al., 2006). In the individual analyses for the reversal learning phase, we first ran an unadjusted model which only included the main factors (i.e. *day, group,* and *sex*) and their interactions for our main behavioral outcome measures (i.e. *probability of choosing the better option, number of rewards, omissions*, and *latencies*) for which a phase interaction was obtained. This was followed by an adjusted model, which included discrimination sessions to criterion as a covariate. The adjusted model was generated to ensure that differences between discrimination and reversal measures were not due to training differences between groups or within individual animals.

Learning data were analyzed with GLM (*fitglme* function; Statistics and Machine Learning Toolbox; MathWorks, Natick, Massachusetts; Version R2017a), with learning phase (discrimination vs reversal), probability group (90-30 vs 70-30), and sex (male vs female) as fixed factors, and individual rat as random factor. All Bonferroni post-hoc tests were corrected for number of comparisons. Statistical significance was noted when p-values were less than 0.05, and p-values between 0.05 and 0.07 were reported as a trend, or marginally significant. Major dependent variables include: probability correct, number of rewards (sucrose pellets earned), total initiation omissions (failure to initiate a trial), total choice omissions (failure to select a stimulus), and median latencies (to initiate a trial, to nosepoke the correct stimulus, to nosepoke the incorrect stimulus, and to collect reward). The latter we refer to as initiation-, correct-, incorrect-, and reward latencies, respectively.

Each trial was classified as a *win* if an animal received a sucrose pellet, and as a *loss* if no reward was delivered. Decisions were classified as better if the animal chose the more rewarding stimulus (stimulus with the larger probability of reward) and *worse* if it chose the less rewarding stimulus. We classified decisions as *Stays* when a rat chose the same stimulus on the subsequent trial and as *Shifts* when it switched to the other alternative. From these first-order measures we were able to construct win-stay, the probability of choosing the same stimulus on the following trial after being rewarded, and lose-shift, the probability of choosing the alternative stimulus after not receiving a reward. These were further parsed into better or worse win-stay or lose-shift, depending on whether the win-stay/lose-shift followed selection of the better option or the worse option. Because we were primarily interested in the differences between the initial phases of discrimination and reversal learning, only the first seven sessions (i.e., early phase learning) for each animal were included in our analysis on response to reward feedback.

#### Metrics of repetition and perseveration

We also used two additional higher-order measures: repetition index and perseveration index. We calculated repetition index (Soltani, Noudoost, & Moore, 2013) as the difference between the actual probability of staying and the chance level of staying:

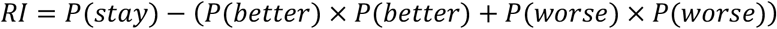

where *P*(*stay*) is the actual probability of staying equal to the joint probability of choosing the option on two consecutive trials, *P*(*better*) is the probability of choosing the more rewarded stimulus, and *P*(*worse*) is the probability of choosing the less rewarded stimulus. We further parsed repetition index into *RI_B_* and *RI_W_*, which accounts for differing tendency to stay on better and worse options:

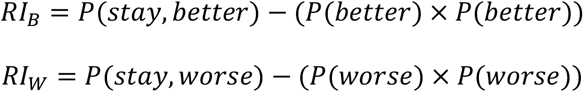

Additionally, we introduced a perseverative index, analogous to the perseverative index defined in Brigman et al. (2008) and Brigman et al. (2010) as the number of same stimulus choices following a loss divided by the number of such “runs,” or the ratio of first-presentation stimulus errors to consecutive errors.

We used these two measures, in addition to the probability of staying, to analyze repetitive behavior. As above, we only used the first seven sessions in calculating repetition measures.

#### Reinforcement learning model

We utilized two simple reinforcement learning models to capture animals’ learning and choice behavior across all sessions during discrimination and reversal. In each model, the estimated subjective reward value of a choice was determined by prior choices and corresponding reward feedback. Specifically, subjective reward values were updated based on reward prediction error, the discrepancy between actual and expected reward. For each observed choice, we updated the subjective reward value of the corresponding option. Accordingly, if the more rewarded stimulus was chosen then *V_choice_* = *V_better_*, and if the less rewarded stimulus was selected *V_choice_* = *V_worse_*, where *V* was the option subjective reward value. We used the following learning rules to update *V_Choice_*:

#### RL1: Model with a single learning rate

On a given trial *t*, the subjective reward value of the chosen stimulus is updated using the following function:

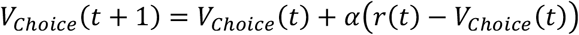

Where *V_Choice_*(*t*) is the subjective reward value on trial *t*, *r*(*t*) is 1 (0) if trial *t* is rewarded (not rewarded) and *α* is the single learning rate.

#### RL2: Model with separate learning rates for rewarded and unrewarded trials

On a trial *t*, the following learning rule was used:

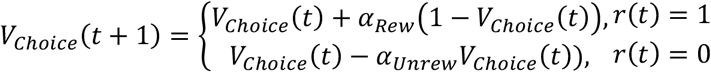

where *α_Rew_* is the learning rate for rewarded trials and *α_Unrew_* is the learning rate for unrewarded trials.

We then applied the following decision rule to determine the probability of selecting the better option on trial *t*:

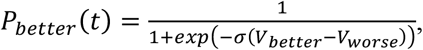

where *σ* is the inverse temperature parameter or sensitivity to the difference in subjective reward values.

We fit each model’s parameters to minimize the negative log likelihood of observed responses. The learning rate parameters (*α*, *α_Rew_*, and *α_Unrew_*) were restricted to values between 0 and 1, and *σ* was constrained to values greater than 0. Learning rate parameters were passed through a sigmoid function to avoid local minima at very low learning rates. Initial parameter values were selected from this range, then fit using MATLAB’s *fmincon* function. For each set of trials, we performed 20 iterations with different initial, randomly selected parameter values to avoid local minima, and the best fit was selected from the iteration with the lowest negative log likelihood (*LL*). Additionally, we also calculated the Bayesian information criterion (BIC) for each fit, defined as

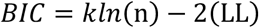

Where *k* is the number of parameters (two in RL1, and three in RL2) and *n* is the number of trials.

We used two methods to examine the relationship between the learning rates and sensitivity to difference in subjective reward values. First, we calculated the Pearson correlation coefficient between estimated learning rate and inverse temperature across animals. That is, after estimating parameters for both groups during discrimination and reversal, we calculated the correlation between sets of learning rates and inverse temperatures across groups during each phase. Second, we examined the parameter correlation derived from our parameter fitting procedure. More specifically, for each animal in each phase, we used the inverse of the Hessian matrix output by *fmincon* to estimate parameter covariance. From this covariance matrix we calculated the correlation matrix. This allowed us to obtain analytic estimates of the correlation between parameters for each animal in each phase (Bishop, 2006; Daw, 2011).

## Results

### Comparisons of Sessions to Criterion, Omnibus measures

#### 90-30 Reward Probability Control Groups

We first conducted statistical analyses to ensure the surgical control groups (s-groups) [i.e., DREADD/VEH, eGFP (CNO+VEH)] that constituted the 90-30 reward probability group did not differ significantly on omnibus measures: number of sessions to criterion (to 70% accuracy), probability of choosing the better option, and number of rewards collected during discrimination and reversal learning (i.e., first seven days of learning common to all rats). There was no effect of surgical control group (GLM: β_s-group_ =4.17, t(5)=0.79, p=0.46) or sex (GLM: β_sex_ =8.17, t(5)=1.55 p=0.18) on sessions to reach criterion, probability correct (GLM: β_s-group_ =0.004, t(54)=0.05, p=0.96; GLM: β_sex_ =0.06, t(54)=0.98, p=0.33), or number of rewards collected (GLM: β_s-group_ =-6.39, t(54)=-0.27, p=0.78; GLM: β_sex_ =2.80, t(54)=0.09, p=0.93) for discrimination learning. Similarly, for reversal learning we did not find any effect of surgical control group (GLM: β_s-group_ =-0.17, t(5)=-0.05, p=0.96) or sex (GLM: β_sex_ =-2.67, t(5)=-0.76, p=0.48) on sessions to reach criterion, probability correct (GLM: β_s-group_ =0.02, t(53)=0.37, p=0.72; GLM: β_sex_ =0.10, t(53)=1.24, p=0.22), or number of rewards collected (GLM: β_s-group_ =-18.52, t(53)=-1.19, p=0.24; GLM: β_sex_ =-7.81, t(53)=-0.40, p=0.69). Given the lack of group differences between surgical control groups within the 90-30 reward probability cohort, we collapsed data across these groups for further analyses.

#### Number of sessions

In total, the 70-30 probability group performed a total of 847 sessions, 417 during the discrimination phase and 430 during reversal (an average of 26.1 ± 3.1 during the discrimination phase and 26.9 ± 1.71 during the reversal phase). The 90-30 probability group performed 270 sessions, 118 during discrimination and 152 during reversal (an average of 13.1 ± 2.6 during discrimination and 16.9 ± 1.65 reversal), **Figure 2B** and **2D**. For discrimination learning, we found no effect of group (p=0.24), sex (p=0.21), or significant group*sex interaction (p=0.49) on sessions to reach criterion. The 90-30 reward probability group required an average of 11.56 ± 2.56 (M ± SEM) sessions while the 70-30 reward probability group required an average of 21.13 ± 3.26 sessions to reach a 70% criterion; females required an average of 11.92 ± 2.47 sessions, while males required an average of 23.92 ± 3.59 sessions. For reversal learning, there was a significant effect of group (GLM: β_group_ =8.28, t(21)=2.63, p=0.02), with the 90-30 reward probability group requiring fewer sessions to reach criterion on average (17.44 ± 1.68) than the 70-30 reward probability (23.50 ± 1.95). However, there was no effect of sex (p=0.20), with males (17.67 ± 1.97) and females (24.69 ± 1.80) requiring a comparable number of sessions to reach criterion for reversal learning, and no significant group*sex interaction (p=0.41). As differing rates of acquisition during discrimination learning could be attributed to differences in performance in the reversal learning phase, discrimination sessions to criterion was included as a covariate in reversal learning analyses (whenever a phase interaction justified analysis of each phase separately), specifically on the main behavioral outcome measures for which those interactions were found. As a preview, the pattern of results were largely consistent with those obtained without the covariate in the model.

**Figure 2.**
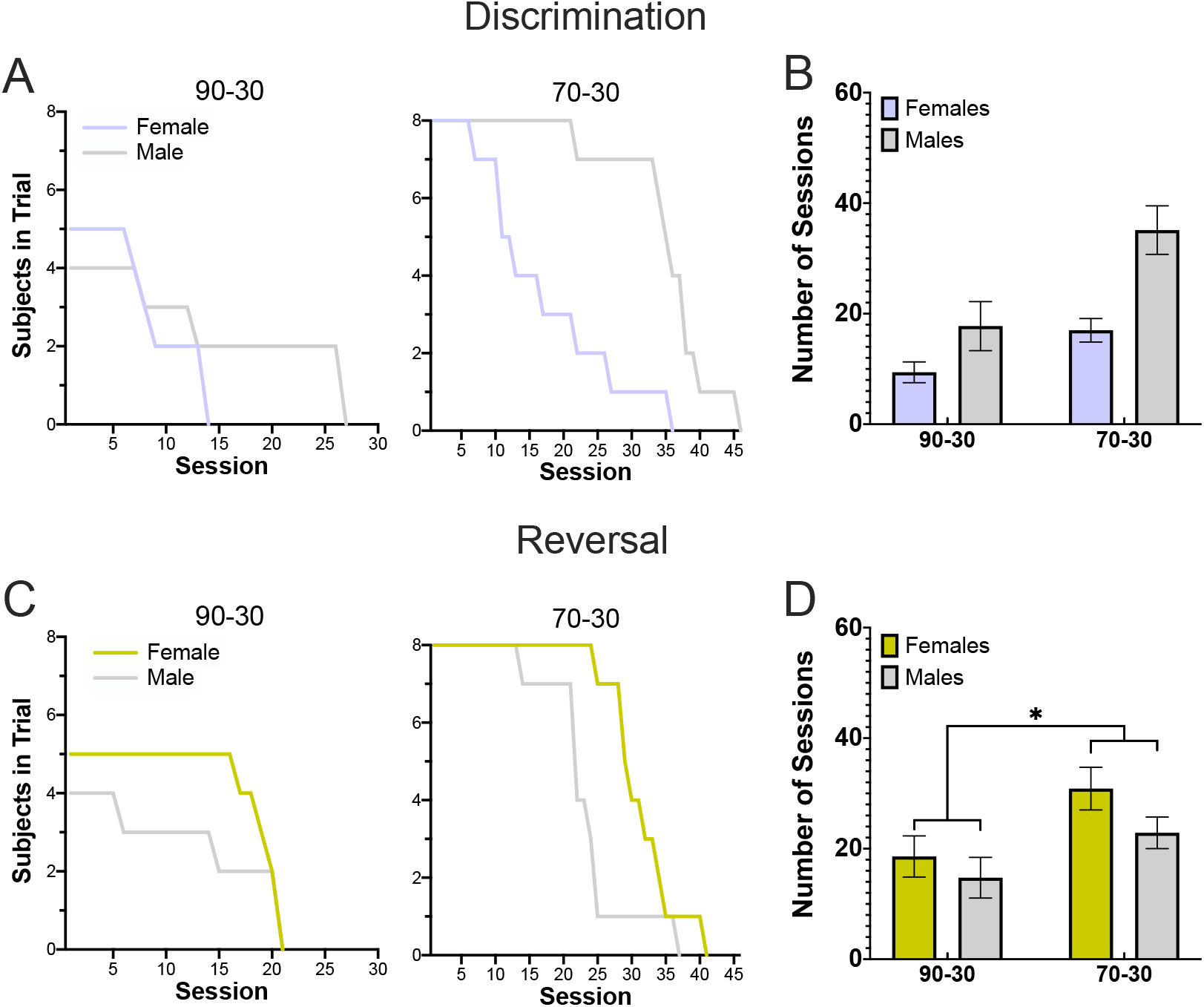
Greater number of completed sessions for the 70-30 reward probability group in both discrimination and reversal learning. **(A-B)** Plotted are the number of subjects per session (A) and the number of sessions to criterion (B) during the discrimination (pre-reversal) phase. The 70-30 reward probability group completes significantly more sessions during discrimination than the 90-30 group. **(C-D)** The same as A-B but during for reversal learning. The 70-30 reward probability group again completes significantly more sessions than the 90-30 group. Bars indicate ± *SEM* *p ≤0.05

### Comparisons of Accuracy and Rewards Collected

#### Comparisons between early Discrimination & Reversal Learning

We fitted GLMs that combined analysis of the first 7 days of learning across both phases (discrimination and reversal). For *probability of choosing the better option* (**Table 1**), we found that all rats exhibited learning by demonstrating an increase in choosing the better option across days (**Figure 3**). Overall animals chose the better option more in the discrimination phase than the reversal phase. For the *number of rewards* (**Table 2**), all rats increased their number of rewards collected across days for both learning phases. There were no difference between reward probability group, or learning phase, nor were there any factor interactions on probability correct or number of rewards. As there were no significant phase interactions, we were not justified to analyze the learning phases separately for these measures.

**Figure 3.**
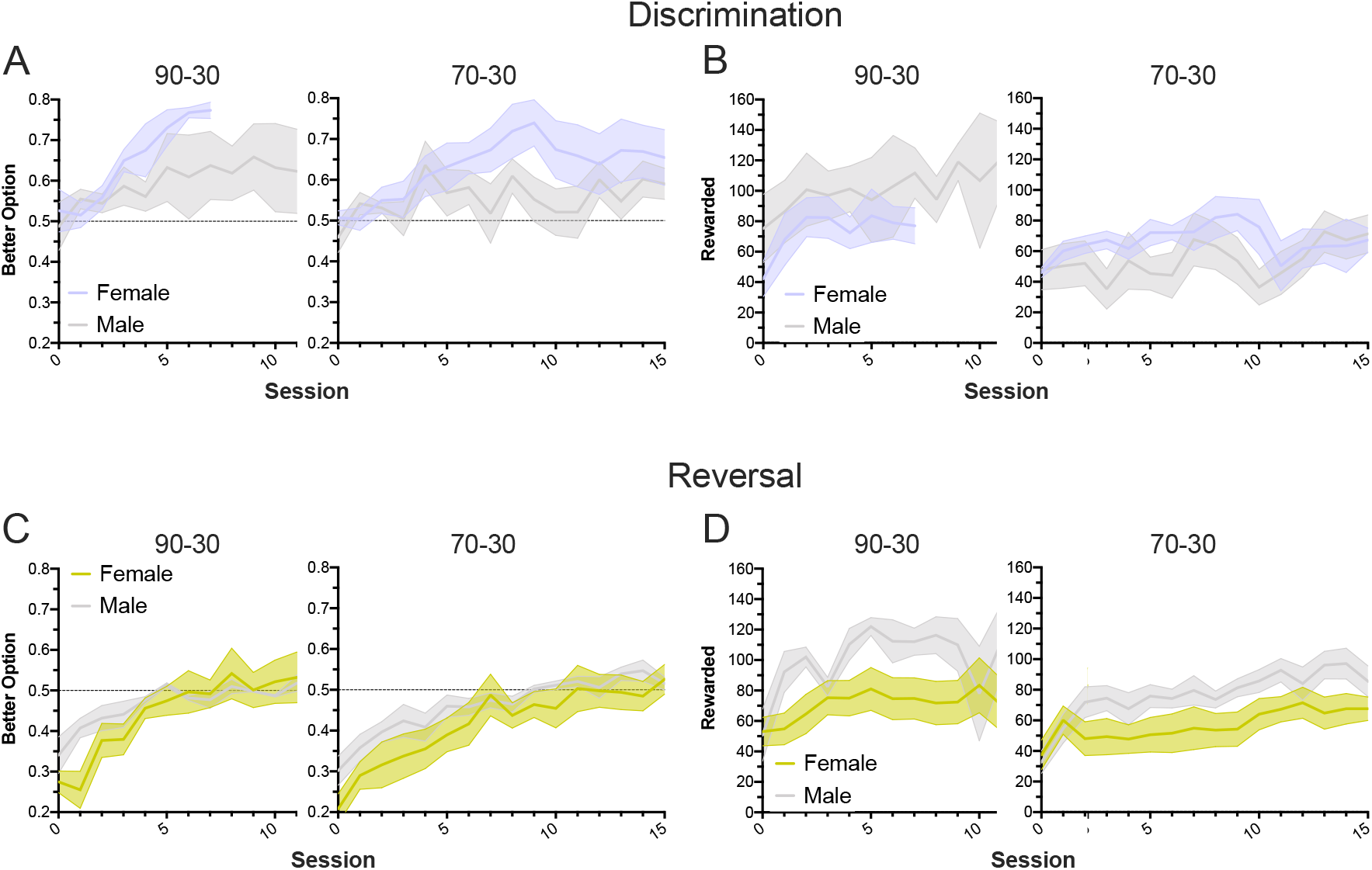
Both reward probability groups and both sexes increase their collected rewards over time but animals choose the better option more often in the discrimination phase. **(A-B)** Proportion of better option selections (A) and number of rewards in a session (B) in the discrimination (pre-reversal) phase, showing the first 10 and 15 trials. Both groups increase selection of the better option and receive more rewards per session over time, with no significant differences between reward probability groups or sex. **(C-D)** Same as A-B, but in the reversal phase. Again, animals in both reward probability groups improve accuracy and collected rewards over time, with no differences by group or sex. Notably, there was significant phase difference on choice of the better option, with the discrimination > reversal phase.

**Table 1.** Probability of choosing the better option during early discrimination and reversal learning

| Early Discrimination and Reversal Learning |  |  |  |  |  |  |  |
| --- | --- | --- | --- | --- | --- | --- | --- |
| $\gamma$ = probability of choosing the better option | | | | | | | |
| Formula | $\gamma \sim [1 + \text{phase} * \text{group} * \text{sex} * \text{day} + (1 \vert \text{rats})]$ | | | | | | |
| Coefficients | $\beta$ | SE | tStat | DF | p | CI<sub>L</sub> | CI<sub>U</sub> |
| intercept | 0.4479 | 0.0463 | 9.6847 | 328 | <.0001 | 0.3570 | 0.5389 |
| **phase** | -0.2293 | 0.0758 | -3.0271 | 328 | **0.0027** | -0.3783 | -0.0803 |
| group | 0.0070 | 0.0535 | 0.1314 | 328 | 0.8955 | -0.0982 | 0.1122 |
| sex | 0.0433 | 0.0763 | 0.5667 | 328 | 0.5713 | -0.1069 | 0.1934 |
| **day** | 0.0462 | 0.0104 | 4.4607 | 328 | **<.0001** | 0.0258 | 0.0666 |
| sex:group | -0.0270 | 0.0848 | -0.3189 | 328 | 0.7500 | -0.1938 | 0.1398 |
| sex:phase | 0.0836 | 0.1000 | 0.8357 | 328 | 0.4039 | -0.1132 | 0.2804 |
| group:phase | -0.0270 | 0.0831 | -0.3251 | 328 | 0.7453 | -0.1905 | 0.1365 |
| sex:day | -0.0270 | 0.0228 | -1.1846 | 328 | 0.2370 | -0.0719 | 0.0179 |
| group:day | -0.0157 | 0.0142 | -1.1069 | 328 | 0.2692 | -0.0435 | 0.0122 |
| phase:day | -0.0040 | 0.0157 | -0.2532 | 328 | 0.8003 | -0.0348 | 0.0269 |
| sex:group:phase | 0.0134 | 0.1138 | 0.1175 | 328 | 0.9065 | -0.2105 | 0.2372 |
| sex:group:day | 0.0152 | 0.0254 | 0.5979 | 328 | 0.5503 | -0.0348 | 0.0652 |
| sex:phase:day | 0.0086 | 0.0247 | 0.3474 | 328 | 0.7285 | -0.0400 | 0.0572 |
| group:phase:day | 0.0070 | 0.0193 | 0.3611 | 328 | 0.7183 | -0.0310 | 0.0449 |

**Table 2.** Total number of rewards collected during early discrimination and reversal learning

| Early Discrimination and Reversal Learning |  |  |  |  |  |  |  |
| --- | --- | --- | --- | --- | --- | --- | --- |
| $\gamma$ =number of rewards | | | | | | | |
| Formula | $\gamma \sim [1 + \text{phase} * \text{group} * \text{sex} * \text{day} + (1 \vert \text{rats})]$ | | | | | | |
| Coefficients | $\beta$ | SE | tStat | DF | p | CI<sub>L</sub> | CI<sub>U</sub> |
| intercept | 50.727 | 10.730 | 4.7278 | 328 | <.0001 | 29.620 | 71.835 |
| phase | -0.6417 | 11.562 | -0.0555 | 328 | 0.9558 | -23.387 | 22.103 |
| group | -2.1959 | 12.206 | -0.1799 | 328 | 0.8573 | -26.207 | 21.815 |
| sex | 29.130 | 17.266 | 1.6871 | 328 | 0.0925 | -4.8371 | 63.097 |
| **day** | 5.9808 | 2.0728 | 2.8854 | 328 | **0.0042** | 1.9031 | 10.058 |
| sex:group | -27.929 | 21.903 | -1.2751 | 328 | 0.2032 | -71.018 | 15.160 |
| sex:phase | -21.182 | 29.708 | -0.7130 | 328 | 0.4764 | -79.624 | 37.260 |
| group:phase | -5.3005 | 15.352 | -0.3453 | 328 | 0.7301 | -35.502 | 24.901 |
| sex:day | -2.4540 | 5.8143 | -0.4221 | 328 | 0.6733 | -13.892 | 8.9840 |
| group:day | -2.1913 | 2.4551 | -0.8925 | 328 | 0.3728 | -7.0211 | 2.6386 |
| phase:day | -1.4236 | 2.2873 | -0.6224 | 328 | 0.5341 | -5.9233 | 3.0760 |
| sex:group:phase | 23.830 | 33.182 | 0.7182 | 328 | 0.4732 | -41.446 | 89.106 |
| sex:group:day | -2.0186 | 6.094 | -0.3312 | 328 | 0.7407 | -14.007 | 9.9698 |
| sex:phase:day | 0.0086 | 0.0247 | 0.3474 | 328 | 0.7285 | -0.0400 | 0.0572 |
| group:phase:day | 0.0070 | 0.0193 | 0.3611 | 328 | 0.7183 | -0.0310 | 0.0449 |

#### Summary

The omnibus comparison across phases of early discrimination and reversal learning revealed that all animals demonstrated an increase in accuracy (i.e. probability of choosing the better option) and an increase in the number of rewards collected across days, regardless of reward probability group or sex. However, animals chose the better option more often in the discrimination phase, compared to reversal as expected.

### Comparisons of Omissions and Latencies

#### Comparisons between early Discrimination & Reversal Learning

Similar to above, we fitted GLMs that combined analysis of the first 7 days of learning across both phases (discrimination and reversal), **Table 3**. For total number of *initiation omissions*, there was a significant phase*group*sex interaction, with post-hoc comparisons revealing 70-30 males exhibited more initiation omissions in the discrimination phase than the reversal phase (p=0.01). There was also a significant group*sex interaction on this measure, but post-hoc tests were not significant after accounting for the number of comparisons. There was no effect of phase, reward probability group, or sex, and no significant phase*group, or phase*sex interactions on this measure. For total number of *choice omissions*, there was no effect of phase, reward probability group, or sex, and no significant phase*group, phase*sex, group*sex, or phase*group*sex interactions.

**Table 3.** Initiation and choice omissions during early discrimination and reversal learning

| Early Discrimination and Reversal Learning |  |  |  |  |  |  |  |
| --- | --- | --- | --- | --- | --- | --- | --- |
| Formula: $\gamma \sim [1 + \text{phase} * \text{group} * \text{sex} + (1 \vert \text{rats})]$ | | | | | | | |
| $\gamma$ =initiation omissions | | | | | | | |
| Coefficients | $\beta$ | SE | tStat | DF | p | CI<sub>L</sub> | CI<sub>U</sub> |
| intercept | 138.40 | 40.087 | 3.4524 | 42 | 0.0013 | 57.500 | 219.30 |
| phase | -18.600 | 17.622 | -1.0555 | 42 | 0.2972 | -54.162 | 16.962 |
| group | -27.900 | 44.500 | -0.6270 | 42 | 0.5341 | -117.71 | 61.906 |
| sex | -58.650 | 51.131 | -1.147 | 42 | 0.2579 | -161.84 | 44.538 |
| phase:group | 69.350 | 48.505 | 1.4298 | 42 | 0.1602 | -28.537 | 167.24 |
| phase:sex | 1.1000 | 28.122 | 0.0391 | 42 | 0.9690 | -55.652 | 57.852 |
| **group:sex** | 164.90 | 70.480 | 2.3397 | 42 | **0.0241** | 22.665 | 307.13 |
| **phase:group:sex** | -158.10 | 61.395 | -2.5751 | 42 | **0.0136** | -282.00 | -34.200 |
| $\gamma$ =choice omissions | | | | | | | |
| Coefficients | $\beta$ | SE | tStat | DF | p | CI<sub>L</sub> | CI<sub>U</sub> |
| intercept | 1.000 | 0.2828 | 3.5355 | 42 | 0.0010 | 0.4292 | 1.5708 |
| phase | 0.200 | 0.3347 | 0.5976 | 42 | 0.5533 | -0.4754 | 0.8754 |
| group | 0.625 | 0.8003 | 0.7809 | 42 | 0.4392 | -0.9902 | 2.2402 |
| sex | 3.750 | 2.7390 | 1.3691 | 42 | 1.3691 | -1.7774 | 9.2774 |
| phase:group | -0.700 | 0.5751 | -1.2172 | 42 | 0.2303 | -1.8606 | 0.4606 |
| phase:sex | -2.450 | 2.0085 | -1.2198 | 42 | 0.2293 | -6.5032 | 1.6032 |
| group:sex | -4.625 | 2.8545 | -1.6202 | 42 | 0.1127 | -10.386 | 1.1357 |
| phase:group:sex | 2.575 | 2.0843 | 1.2354 | 42 | 0.2235 | -1.6313 | 6.7813 |

Next we analyzed median initiation and better choice latencies (**Table 4**), as well as worse choice and reward collection latencies (**Table 5)**. For *initiation latencies*, we found a marginal group*sex interaction, but no effect phase, or reward probability group, and no significant phase*group, phase*sex, group*sex, or phase*group*sex interactions. For *better choice latencies*, there was a significant phase*sex interaction, with males exhibiting longer choice latencies for the better option in the discrimination phase compared to the reversal phase (p=0.01), but no effect of phase, reward probability group, or sex, and no significant phase*group, group*sex, or phase*group*sex interactions. For *worse choice latencies*, there was a significant effect of phase, with animals in the reversal phase exhibiting longer latencies than in the discrimination phase, but no effect of reward probability group, sex, or significant phase*group, phase*sex, group*sex, phase*group*sex interactions. For *reward collection latencies*, we found a significant effect of reward probability group, with the 90-30 group taking longer to collect reward than the 70-30 group, but no effect of phase or sex. There was a significant phase*sex interaction, with females exhibiting longer latencies than males in the reversal phase (p=0.01), and a phase*group*sex interaction, with 90-30 males in the discrimination phase than 90-30 males in the reversal phase, but no significant phase*group or group*sex interaction.

**Table 4.** Initiation and better choice latency during early discrimination and reversal learning

| Early Discrimination and Reversal Learning |  |  |  |  |  |  |  |
| --- | --- | --- | --- | --- | --- | --- | --- |
| Formula: $\gamma \sim [1 + \text{phase} * \text{group} * \text{sex} + (1 \vert \text{rats})]$ | | | | | | | |
| $\gamma$=initiation latency | | | | | | | |
| Coefficients | $\beta$ | SE | tStat | DF | p | CI<sub>L</sub> | CI<sub>U</sub> |
| intercept | 4.5118 | 0.8013 | 5.6307 | 42 | <.0001 | 2.8947 | 6.1289 |
| phase | -0.8195 | 0.4883 | -1.6782 | 42 | 0.1007 | -1.8049 | 0.1660 |
| group | -0.2730 | 0.9112 | -0.2996 | 42 | 0.7660 | -2.1118 | 1.5658 |
| sex | -1.1703 | 1.0509 | -1.1136 | 42 | 0.2718 | -3.2911 | 0.9505 |
| phase:group | 1.0286 | 0.9268 | 1.1098 | 42 | 0.2734 | -0.8418 | 2.8991 |
| phase:sex | -0.5760 | 0.9377 | -0.6143 | 42 | 0.5423 | -2.4683 | 1.3163 |
| group:sex | 3.3462 | 1.7256 | 1.9392 | 42 | 0.0592 | -0.1362 | 6.8287 |
| phase:group:sex | -2.4249 | 1.7569 | -1.3802 | 42 | 0.1748 | -5.9706 | 1.1207 |
| $\gamma$=better choice latency | | | | | | | |
| Coefficients | $\beta$ | SE | tStat | DF | p | CI<sub>L</sub> | CI<sub>U</sub> |
| intercept | 0.9025 | 0.0704 | 12.818 | 42 | <.0001 | 0.7604 | 1.0446 |
| phase | 0.0332 | 0.0737 | 0.4503 | 42 | 0.6548 | -0.1156 | 0.1820 |
| group | 0.1099 | 0.1540 | 0.7138 | 42 | 0.4793 | -0.2009 | 0.4208 |
| sex | -0.1401 | 0.1273 | -1.1005 | 42 | 0.2774 | -0.3971 | 0.1169 |
| phase:group | -0.2615 | 0.1338 | -1.9537 | 42 | 0.0574 | -0.5315 | 0.0086 |
| **phase:sex** | -0.2118 | 0.0785 | -2.6991 | 42 | **0.0100** | -0.3702 | -0.0534 |
| group:sex | -0.0796 | 0.2092 | -0.3804 | 42 | 0.7056 | -0.5017 | 0.3426 |
| phase:group:sex | 0.2999 | 0.1648 | 1.8193 | 42 | 0.0760 | -0.0328 | 0.6325 |

**Table 5.** Worse choice and reward latency during early discrimination and reversal learning

| Early Discrimination and Reversal Learning |  |  |  |  |  |  |  |
| --- | --- | --- | --- | --- | --- | --- | --- |
| Formula: $\gamma \sim [1 + \text{phase} * \text{group} * \text{sex} + (1 \vert \text{rats})]$ | | | | | | | |
| $\gamma$ =worse choice latency | | | | | | | |
| Coefficients | $\beta$ | SE | tStat | DF | p | CI<sub>L</sub> | CI<sub>U</sub> |
| intercept | 0.8044 | 0.0824 | 9.7582 | 42 | <.0001 | 0.6380 | 0.9708 |
| **phase** | 0.0965 | 0.0312 | 3.0977 | 42 | **0.0035** | 0.0336 | 0.1594 |
| group | 0.0885 | 0.1358 | 0.6519 | 42 | 0.5181 | -0.1856 | 0.3627 |
| sex | -0.1177 | 0.1553 | -0.7575 | 42 | 0.4530 | -0.4311 | 0.1958 |
| phase:group | -0.1059 | 0.0919 | -1.1521 | 42 | 0.2558 | -0.2913 | 0.0796 |
| phase:sex | -0.1543 | 0.0828 | -1.8636 | 42 | 0.0694 | -0.3213 | 0.0128 |
| group:sex | -0.0308 | 0.2051 | -0.1502 | 42 | 0.8814 | -0.4446 | 0.3830 |
| phase:group:sex | 0.0965 | 0.1379 | 0.6999 | 42 | 0.4879 | -0.1818 | 0.3748 |
| $\gamma$ =reward collection latency | | | | | | | |
| Coefficients | $\beta$ | SE | tStat | DF | p | CI<sub>L</sub> | CI<sub>U</sub> |
| intercept | 1.9392 | 0.1232 | 15.737 | 42 | <.0001 | 1.6905 | 2.1879 |
| phase | -0.0482 | 0.0677 | -0.7120 | 42 | 0.4804 | -0.1848 | 0.0884 |
| **group** | -0.5845 | 0.1387 | -4.2155 | 42 | **0.0001** | -0.8643 | -0.3047 |
| sex | -0.1688 | 0.2034 | -0.8300 | 42 | 0.4113 | -0.5793 | 0.2417 |
| phase:group | 0.0313 | 0.0781 | 0.4001 | 42 | 0.6911 | -0.1264 | 0.1890 |
| **phase:sex** | -0.2257 | 0.0906 | -2.4909 | 42 | **0.0168** | -0.4085 | -0.0428 |
| group:sex | 0.2706 | 0.2202 | 1.2288 | 42 | 0.2260 | -0.1738 | 0.7151 |
| **phase:group:sex** | 0.2924 | 0.1188 | 2.4604 | 42 | **0.0181** | 0.0526 | 0.5322 |

#### Comparisons within Discrimination Learning

Because we found phase interactions on initiation omissions, better choice latencies, and reward collection latencies, we used GLMs to analyze these variables for the discrimination phase separately (**Table 6**). For total number of *initiation omissions*, there was no significant effects of reward probability group, sex, or group*sex interaction (**Figure 4A**). For *better choice latencies*, there was also no effect of reward probability group or sex, or group*sex interaction, despite a phase by sex interaction found during early learning (**Figures 4B, S1A, and S1B**). However, for *reward collection latencies*, there was a significant effect of reward probability group with the 90-30 group taking longer to collect reward than the 70-30 group, but no effect of sex and no group*sex interaction (**Figures 4C, S1C, and S1D**).

**Figure 4.**
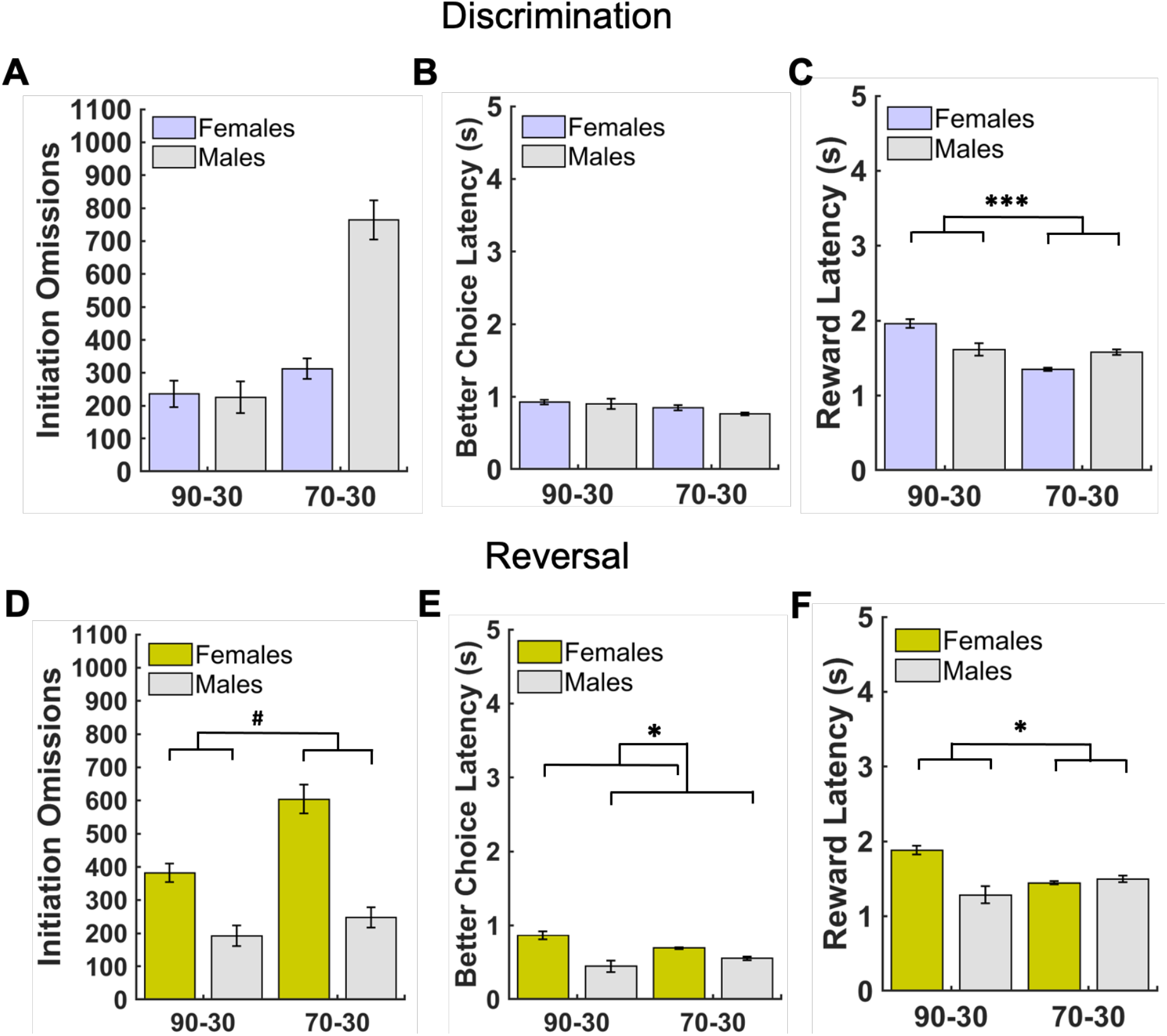
Patterns of latencies by sex and reward probability group during discrimination and reversal learning. (A) There were no group or sex differences in initation omissions in discrimination. (B) There were no group or sex differences in better choice latencies in discrimination. (C) There were significant probability group differences in reward collection latencies in discrimination, with the 90-30 reward probability group exhibiting longer latencies than the 70-30 reward probability group. (D) There were no group or sex differences in initation omissions in reversal. (E) There were sex differences in better choice latencies in the reversal phase, with females taking longer to make a choice of the better option than males (with and without controlling for the number discrimination sessions to criterion). (F) There were significant probability group differences in reward collection latencies in the reversal phase, with the 90-30 reward probability group exhibiting longer latencies than the 70-30 reward probability group (with and without controlling for the number of discrimination sessions to criterion). Bars indicate ± SEM ^#^p=0.07, *p ≤0.05, ***p≤0.001

**Table 6.** Initiation omissions, better choice latency, and reward collection latency during discrimination learning

| Discrimination Learning |  |  |  |  |  |  |  |
| --- | --- | --- | --- | --- | --- | --- | --- |
| Formula: $\gamma \sim [1 + \text{group} * \text{sex}]$ | | | | | | | |
| $\gamma$ =initiation omissions | | | | | | | |
| Coefficients | $\beta$ | SE | tStat | DF | p | CI<sub>L</sub> | CI<sub>U</sub> |
| intercept | 236.00 | 145.72 | 1.6195 | 21 | 0.1203 | -67.05 | 539.05 |
| group | 76.125 | 185.76 | 0.4098 | 21 | 0.6861 | -310.19 | 462.44 |
| sex | -11.000 | 218.59 | -0.0503 | 21 | 0.9603 | -465.58 | 443.58 |
| group:sex | 462.88 | 272.63 | 1.6978 | 21 | 0.1043 | -104.08 | 1029.8 |
| $\gamma$ =better choice latency | | | | | | | |
| Coefficients | $\beta$ | SE | tStat | DF | p | CI<sub>L</sub> | CI<sub>U</sub> |
| intercept | 0.9173 | 0.1052 | 8.7342 | 21 | <.0001 | 0.6989 | 1.1357 |
| group | 0.0555 | 0.1339 | 0.4147 | 21 | 0.6826 | -0.2229 | 0.3339 |
| sex | -0.0619 | 0.1575 | -0.3931 | 21 | 0.6982 | -0.3895 | 0.2657 |
| group:sex | -0.1631 | 0.1965 | -0.8303 | 21 | 0.4157 | -0.5717 | 0.2455 |
| $\gamma$ =reward collection latency | | | | | | | |
| Coefficients | $\beta$ | SE | tStat | DF | p | CI<sub>L</sub> | CI<sub>U</sub> |
| intercept | 1.9705 | 0.1200 | 16.42 | 21 | <.0001 | 1.7209 | 2.2201 |
| **group** | -0.6006 | 0.1530 | -3.9258 | 21 | **0.0008** | -0.9187 | -0.2824 |
| sex | -0.2429 | 0.1800 | -1.3492 | 21 | 0.1916 | -0.6172 | 0.1315 |
| group:sex | 0.3960 | 0.2245 | 1.7638 | 21 | 0.0923 | -0.0709 | 0.8629 |

#### Comparisons within Reversal Learning

As above, because we found phase interactions on initiation omissions, better choice latencies, and reward collection latencies, we used GLMs to analyze these variables for the reversal phase separately. We ran two models for the reversal learning phase: an unadjusted model which included only the main factors (i.e. *group* and *sex*), and an adjusted model with the number of discrimination sessions to criterion added as a covariate (**Tables 7 and 8**). For total number of *initiation omissions*, we found a marginal effect of reward probability group (p=0.07) in the adjusted model, but did not find a significant sex, or group*sex interaction with either model (**Figure 4D)**. For *better choice latencies*, there was an effect of sex in both the unadjusted and adjusted model, with females exhibiting longer choice latencies than males when choosing the better option. No significant group or group*sex interactions were found for either model (**Figures 4E, S1E, and S1F**). For *reward collection latencies*, there was a significant effect of reward probability group, with the 90-30 group taking longer to collect reward than the 70-30 group, in both the unadjusted and adjusted model, but no significant sex or group*sex interaction was observed (**Figures 4F, S1G and S1H**).

**Table 7.** Initiation omissions, better choice latency, and reward collection latency during reversal learning

| Reversal Learning |  |  |  |  |  |  |  |
| --- | --- | --- | --- | --- | --- | --- | --- |
| Formula: $\gamma \sim [1 + \text{group} * \text{sex}]$ | | | | | | | |
| $\gamma$ =initiation omissions | | | | | | | |
| Coefficients | $\beta$ | SE | tStat | DF | p | CI<sub>L</sub> | CI<sub>U</sub> |
| intercept | 381.80 | 113.35 | 3.3682 | 21 | 0.0029 | 146.07 | 617.53 |
| group | 222.08 | 144.50 | 1.5369 | 21 | 0.1393 | -78.426 | 522.58 |
| sex | -189.30 | 170.03 | -1.1133 | 21 | 0.2782 | -542.90 | 164.30 |
| group:sex | -167.08 | 212.07 | -0.7878 | 21 | 0.4396 | -608.09 | 273.94 |
| $\gamma$ =better choice latency | | | | | | | |
| Coefficients | $\beta$ | SE | tStat | DF | p | CI<sub>L</sub> | CI<sub>U</sub> |
| intercept | 0.8899 | 0.0930 | 9.5727 | 21 | <.0001 | 0.6966 | 1.0832 |
| group | -0.1766 | 0.1185 | -1.4901 | 21 | 0.1511 | -0.4230 | 0.0699 |
| **sex** | -0.3229 | 0.1394 | -2.3156 | 21 | **0.0308** | -0.6129 | -0.0329 |
| group:sex | 0.2153 | 0.1739 | 1.2382 | 21 | 0.2293 | -0.1463 | 0.5770 |
| $\gamma$ =reward collection latency | | | | | | | |
| Coefficients | $\beta$ | SE | tStat | DF | p | CI<sub>L</sub> | CI<sub>U</sub> |
| Intercept | 1.8941 | 0.1442 | 13.139 | 21 | <.0001 | 1.5943 | 2.1939 |
| **group** | -0.4881 | 0.1838 | -2.6560 | 21 | **0.0148** | -0.8703 | -0.1059 |
| sex | -0.3879 | 0.2163 | -1.7936 | 21 | 0.0873 | -0.8376 | 0.0619 |
| group:sex | 0.4887 | 0.2697 | 1.8121 | 21 | 0.0843 | -0.0722 | 1.0496 |

**Table 8.** Initiation omissions, better choice latency, and reward collection latency during reversal learning with Discrimination Sessions to Criterion (dis_stc) as a covariate

| Reversal Learning (Covariate: Discrimination Sessions to Criterion) |  |  |  |  |  |  |  |
| --- | --- | --- | --- | --- | --- | --- | --- |
| Formula: $\gamma \sim [1 + \text{group} * \text{sex} + \text{dis\_stc}]$ | | | | | | | |
| $\gamma$ = initiation omissions | | | | | | | |
| Coefficients | $\beta$ | SE | tStat | DF | p | CIL | CIU |
| intercept | 445.62 | 116.21 | 3.8344 | 20 | 0.0010 | 203.2 | 688.04 |
| group | 272.93 | 142.18 | 1.9197 | 20 | 0.0693 | -23.643 | 569.50 |
| sex | -125.48 | 167.93 | -0.7472 | 20 | 0.4636 | -475.77 | 224.81 |
| dis_stc | -7.9773 | 5.2301 | -1.5253 | 20 | 0.1428 | -18.887 | 2.9325 |
| group:sex | -123.20 | 204.87 | -0.6014 | 20 | 0.5544 | -550.55 | 304.15 |
| $\gamma$ = better choice latency | | | | | | | |
| Coefficients | $\beta$ | SE | tStat | DF | p | CIL | CIU |
| intercept | 0.8958 | 0.0996 | 8.9951 | 20 | <.0001 | 0.6881 | 1.1036 |
| group | -0.1719 | 0.1218 | -1.4106 | 20 | 0.1737 | -0.4260 | 0.0823 |
| **sex** | -0.3170 | 0.1439 | -2.2026 | 20 | **0.0395** | -0.6172 | -0.0168 |
| dis_stc | -0.0007 | 0.0045 | -0.1653 | 20 | 0.8570 | -0.0101 | 0.0086 |
| group:sex | 0.2194 | 0.1756 | 1.2498 | 20 | 0.2258 | 0.1468 | 0.5856 |
| $\gamma$ = reward collection latency | | | | | | | |
| Coefficients | $\beta$ | SE | tStat | DF | p | CIL | CIU |
| intercept | 1.8448 | 0.1521 | 12.131 | 20 | <.0001 | 1.5275 | 2.162 |
| **group** | -0.5274 | 0.1861 | -2.8348 | 20 | **0.0102** | -0.9155 | -0.1393 |
| sex | -0.4372 | 0.2198 | -1.9894 | 20 | 0.0605 | -0.8956 | 0.0212 |
| dis_stc | 0.0062 | 0.0068 | 0.9009 | 20 | 0.3784 | -0.0081 | 0.0204 |
| group:sex | 0.4548 | 0.2681 | 1.6965 | 20 | 0.1053 | -0.1044 | 1.014 |

#### Summary

After controlling for the number of discrimination sessions to criterion in reversal learning, we observed the same pattern of effects as that obtained with the original model: female animals exhibited longer choice latencies for the better option than males (this pattern was not observed for the discrimination phase). Additionally, we found longer reward collection latencies in animals learning the 90-30 reward probabilities, compared to animals in the 70-30 group. This effect was observed in both the discrimination and reversal phases. Finally, though we initially found a phase interaction for initiation omissions in early learning, follow-up analyses yielded only a marginal effect of group, but no sex, or group by sex interactions when the phases were analyzed separately. In sum, we determined that latencies, and not omissions, are among the more sensitive measures of performance; certainly beyond omnibus measures of accuracy and cached rewards that are typically reported in the literature.

### Comparisons of Response to Reward Feedback

#### Comparisons between early Discrimination & Reversal Learning

We analyzed win-stay and lose-shift strategies during early discrimination and reversal learning (**Table 9**). For win-stay, we found a marginally significant effect of phase on employing the win-stay strategy, with animals using this strategy more in the discrimination phase than in the reversal phase, a significant effect of reward probability group, with animals in the 90-30 reward probability group using this strategy more than those in the 70-30 reward probability group, but no effect of sex. There was also a significant phase*group interaction, with animals in the 90-30 reward probability group using this strategy more in the discrimination phase than the reversal phase (p=0.001), but no significant phase*sex, group*sex, or phase*group*sex interactions. For lose-shift, there was a significant effect of phase, with animals employing this strategy more in the discrimination learning phase than the reversal phase, a significant effect of reward probability group, with the 90-30 group using the lose-shift strategy more than the 70-30 group. Finally, there was also an effect of sex, with males employing this strategy more than females, but no significant phase*group, phase*sex, group*sex, phase*group*sex interactions.

**Table 9.** Win-stay and lose-shift during early discrimination and reversal learning

| Early Discrimination and Reversal Learning |  |  |  |  |  |  |  |
| --- | --- | --- | --- | --- | --- | --- | --- |
| Formula: $\gamma \sim [1 + \text{phase} * \text{group} * \text{sex} + (1 \vert \text{rats})]$ | | | | | | | |
| $\gamma = \text{win-stay}$ | | | | | | | |
| Coefficients | $\beta$ | SE | tStat | DF | p | CI<sub>L</sub> | CI<sub>U</sub> |
| intercept | 0.5941 | 0.0122 | 48.508 | 42 | <.0001 | 0.5694 | 0.6188 |
| **phase** | -0.1269 | 0.0290 | -4.3795 | 42 | **0.0001** | -0.1853 | -0.0684 |
| **group** | -0.0746 | 0.0344 | -2.1708 | 42 | **0.0357** | -0.1440 | -0.0052 |
| sex | -0.0876 | 0.0551 | -1.5900 | 42 | 0.1193 | -0.1988 | 0.0236 |
| **phase:group** | 0.1068 | 0.0512 | 2.0834 | 42 | **0.0433** | 0.0033 | 0.2102 |
| phase:sex | 0.0442 | 0.0621 | 0.7118 | 42 | 0.4805 | -0.0811 | 0.1695 |
| group:sex | 0.0403 | 0.0667 | 0.6036 | 42 | 0.5494 | -0.0944 | 0.1750 |
| phase:group:sex | -0.0328 | 0.0782 | -0.4196 | 42 | 0.6769 | -0.1907 | 0.1251 |
| $\gamma = \text{lose-switch}$ | | | | | | | |
| Coefficients | $\beta$ | SE | tStat | DF | p | CI<sub>L</sub> | CI<sub>U</sub> |
| intercept | 0.5721 | 0.0156 | 36.586 | 42 | <.0001 | 0.5405 | 0.6036 |
| **phase** | -0.1572 | 0.0370 | -4.2478 | 42 | **0.0001** | -0.2318 | -0.0825 |
| **group** | -0.0634 | 0.0192 | -3.3082 | 42 | **0.0019** | -0.1020 | -0.0247 |
| **sex** | 0.0398 | 0.0186 | 2.1392 | 42 | **0.0383** | 0.0023 | 0.0773 |
| phase:group | 0.0345 | 0.0487 | 0.7082 | 42 | 0.4827 | -0.0638 | 0.1328 |
| phase:sex | 0.0325 | 0.0409 | 0.7942 | 42 | 0.4316 | -0.0501 | 0.1150 |
| group:sex | -0.0321 | 0.0253 | -1.2661 | 42 | 0.2124 | -0.0832 | 0.0191 |
| phase:group:sex | 0.0368 | 0.0561 | 0.6564 | 42 | 0.5152 | -0.0763 | 0.1499 |

#### Comparisons within Discrimination Learning

Because we found phase interactions on win-stay, we were justified to analyze potential differences in win-stay strategies on stimulus responses employed by each reward probability group and by sex, in the discrimination phase separately (**Table 10)**. For win-stay, there was an effect of reward probability group, but no effect of sex, and no group*sex interaction (**Figure 5A**). When we considered win-stay *on the better option* (win-stay|better), we found a marginally significant effect of reward probability group, with the 90-30 group employing this strategy more, but no effect of sex or significant group*sex interaction (**Figure 5B**). When we analyzed win-stay on the worse option (win-stay|worse), we found no significant effect of reward probability group, sex, or group*sex interaction (**Figure 5C**).

**Figure 5.**
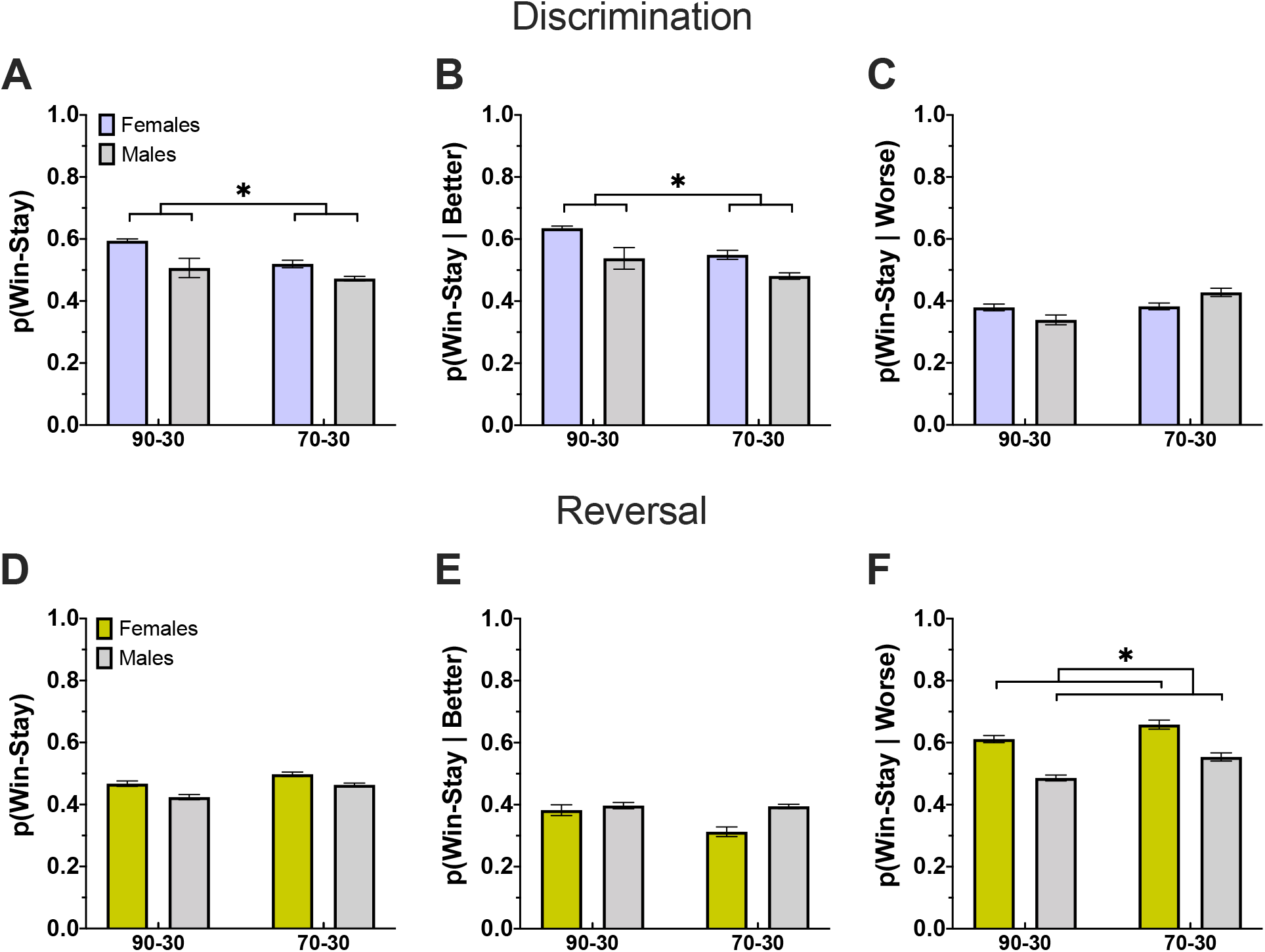
Greater overall win-stay and win-stay on the better option in the 90-30 group during discrimination. (**A-C)** Plotted are proportion of win-stay responses overall (A), after choosing the better option (B), and after choosing the worse option (C) during discrimination. Overall win-stay and win-stay on the better option is used more often in the 90-30 group than the 70-30 group. **(D-F)** The same as A-C but for reversal. We find no significant effects on overall win-stay and win-stay on the better option, but do find females are more likely to apply a win-stay strategy after choosing the worse option than males. Bars indicate ± *SEM* *p ≤0.05.

**Table 10.**
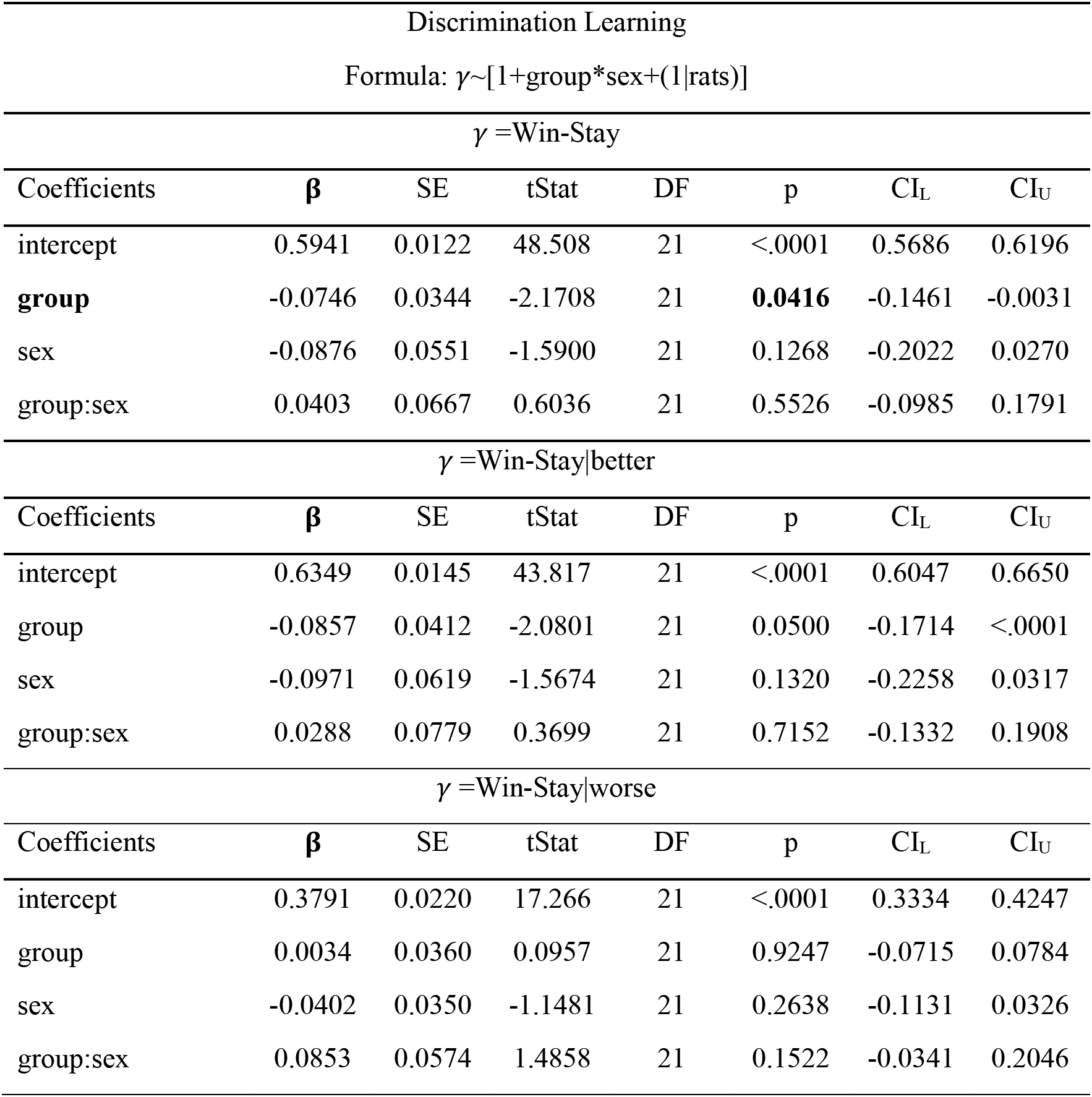
Win-stay, win-stay|better, and win-stay|worse during discrimination learning

#### Comparisons within Reversal Learning

Per the phase interactions, we were permitted to analyze win-stay (**Table 11**) strategies on stimulus responses employed during the reversal phase, separately. We found males were less likely to employ a win-stay strategy (**Figure 5D)**, but no effect of reward probability group, or group*sex interaction. We found no significant group or sex differences, and no group*sex interaction for win-stay|better (**Figure 5E**). We did, however, find a significant effect of sex on win-stay|worse, with males less likely to employ the strategy than females, but no reward probability group effect or group*sex interaction (**Figure 5F**).

**Table 11.**
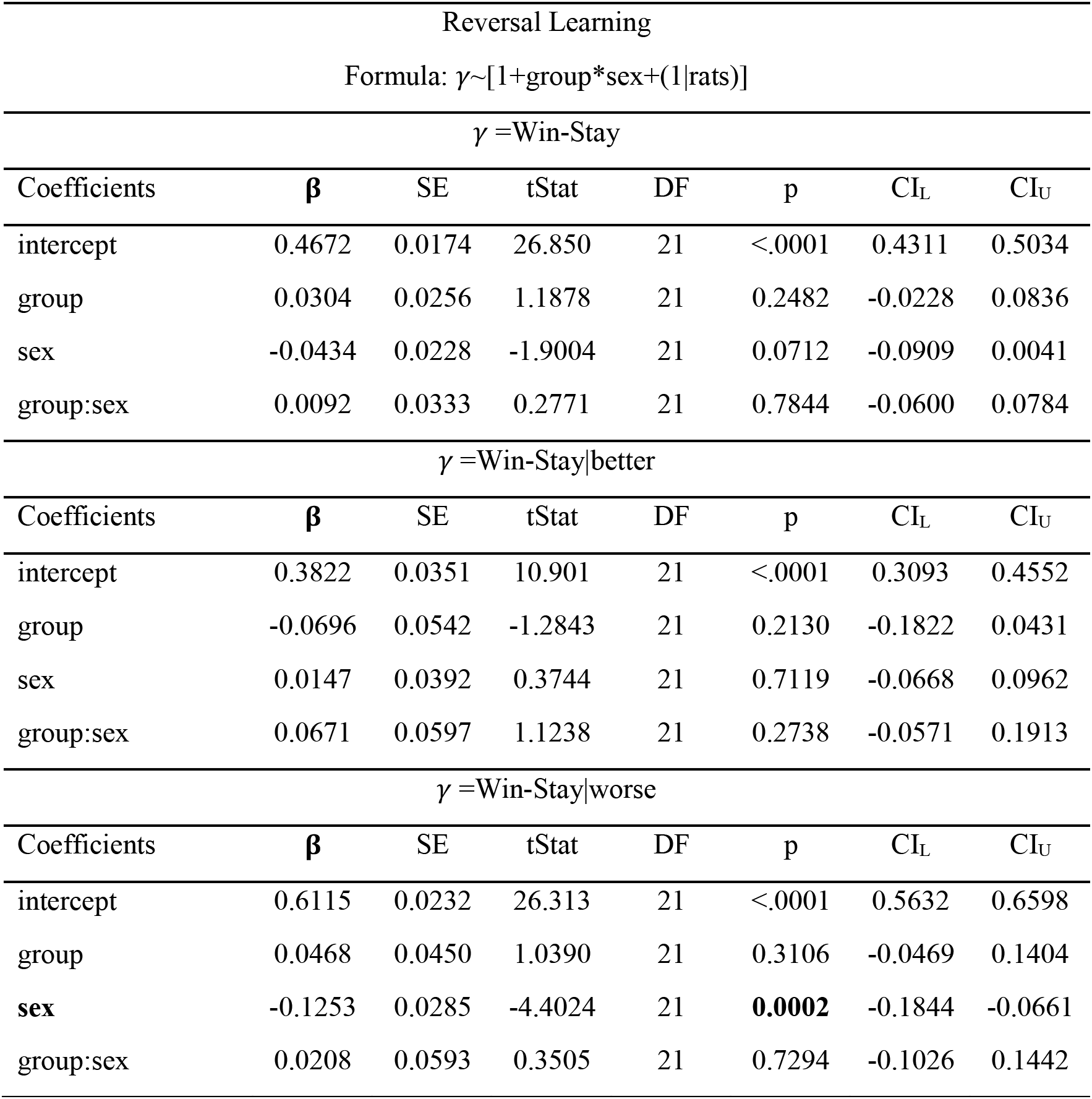
Win-stay, win-stay|better, and win-stay|worse during reversal learning

| Reversal Learning |  |  |  |  |  |  |  |
| --- | --- | --- | --- | --- | --- | --- | --- |
| Formula: $\gamma \sim [1 + \text{group} * \text{sex} + (1 \vert \text{rats})]$ | | | | | | | |
| $\gamma = \text{Win-Stay}$ | | | | | | | |
| Coefficients | $\beta$ | SE | tStat | DF | p | CI<sub>L</sub> | CI<sub>U</sub> |
| intercept | 0.4672 | 0.0174 | 26.850 | 21 | <.0001 | 0.4311 | 0.5034 |
| group | 0.0304 | 0.0256 | 1.1878 | 21 | 0.2482 | -0.0228 | 0.0836 |
| sex | -0.0434 | 0.0228 | -1.9004 | 21 | 0.0712 | -0.0909 | 0.0041 |
| group:sex | 0.0092 | 0.0333 | 0.2771 | 21 | 0.7844 | -0.0600 | 0.0784 |
| $\gamma = \text{Win-Stay\vert better}$ | | | | | | | |
| Coefficients | $\beta$ | SE | tStat | DF | p | CI<sub>L</sub> | CI<sub>U</sub> |
| intercept | 0.3822 | 0.0351 | 10.901 | 21 | <.0001 | 0.3093 | 0.4552 |
| group | -0.0696 | 0.0542 | -1.2843 | 21 | 0.2130 | -0.1822 | 0.0431 |
| sex | 0.0147 | 0.0392 | 0.3744 | 21 | 0.7119 | -0.0668 | 0.0962 |
| group:sex | 0.0671 | 0.0597 | 1.1238 | 21 | 0.2738 | -0.0571 | 0.1913 |
| $\gamma = \text{Win-Stay\vert worse}$ | | | | | | | |
| Coefficients | $\beta$ | SE | tStat | DF | p | CI<sub>L</sub> | CI<sub>U</sub> |
| intercept | 0.6115 | 0.0232 | 26.313 | 21 | <.0001 | 0.5632 | 0.6598 |
| group | 0.0468 | 0.0450 | 1.0390 | 21 | 0.3106 | -0.0469 | 0.1404 |
| **sex** | -0.1253 | 0.0285 | -4.4024 | 21 | **0.0002** | -0.1844 | -0.0661 |
| group:sex | 0.0208 | 0.0593 | 0.3505 | 21 | 0.7294 | -0.1026 | 0.1442 |

#### Summary

Win-stay and win-stay|better strategies were employed more often during early discrimination learning compared to early reversal learning. Further, the 90-30 reward probability group used both win-stay and win-stay|better strategies more in discrimination learning than the 70-30 group. Lastly, female animals were more likely to apply a win-stay|worse strategy in the reversal phase compared to males.

### Comparisons of Repetition measures

#### Comparisons between early Discrimination & Reversal Learning

We next fitted GLMs to examine the effects of different factors on repetition in choice behavior. We did not find any significant effect of phase, group, sex, or any significant interaction of those factors on overall *p*(*Stay*), p(Stay | better), or p(*Stay* | *worse*) (**Table 12**). However, there was an increased perseveration index in the reversal phase relative to discrimination (**Figures 6A and 6E**), regardless of reward probability group or sex (**Table 13**). For *RI,* we found a significantly lower value in the 70-30 group and in males, but no significant effects of phase, phase*group, phase*sex, group*sex, or phase*group*sex interaction (**Figures 6B and 6F**). Similarly, for *RI_B_* (**Table 14**) we observed lower values in the 70-30 group and in males, but no significant difference by phase, phase*group, phase*sex, group*sex, or phase*group*sex interaction (**Figures 6C and 6G).** This pattern holds for *RI_W_* as well, with a lower *RI_W_* in the 70-30 group and males, and no significant effects of phase, phase*group, phase*sex, group*sex, or group*phase*sex interaction (**Figures 6D and 6H**). As we observed no phase interactions, we were not justified in further analyses of discrimination and reversal phases, separately.

**Figure 6.**
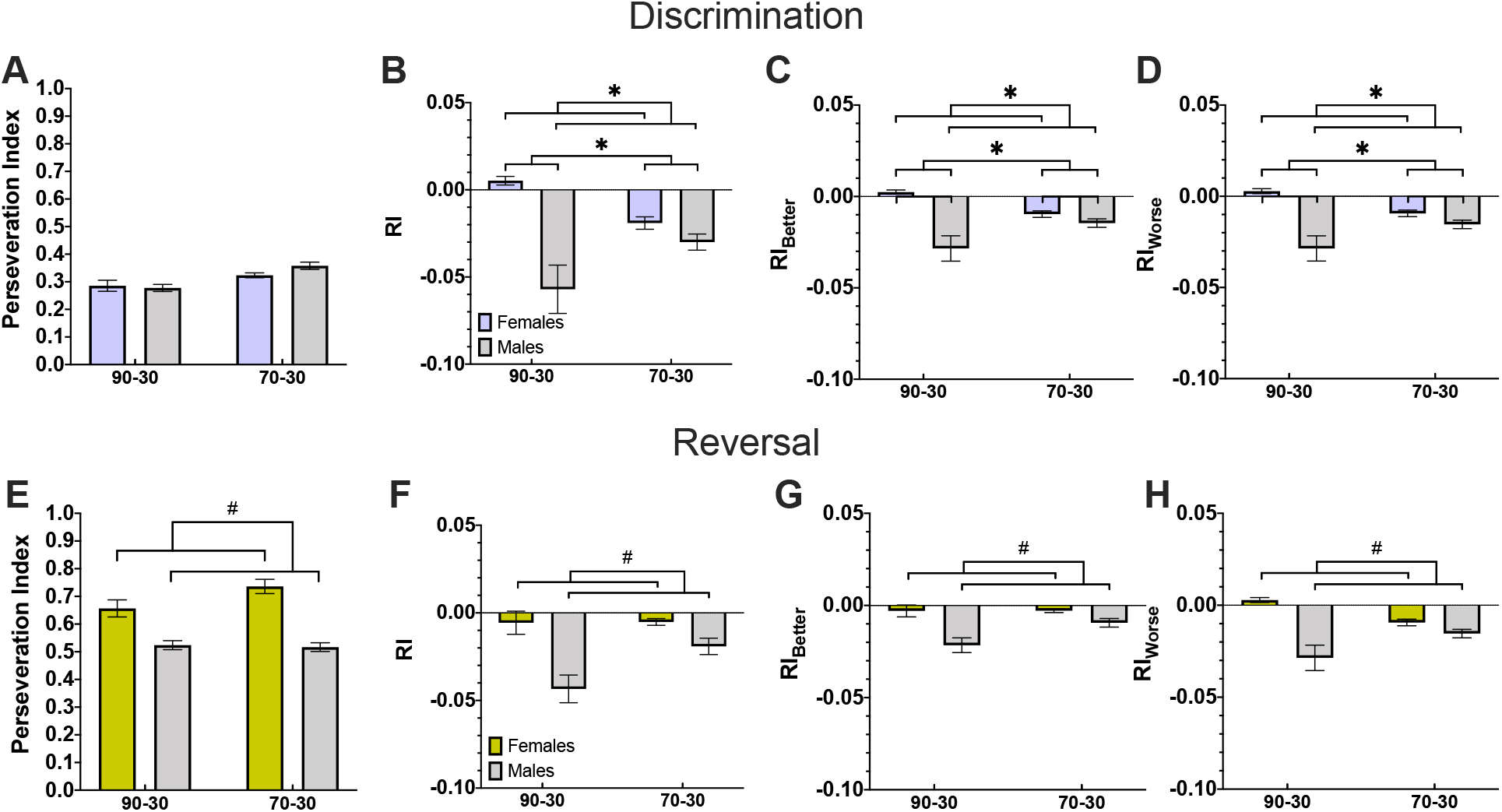
Greater perseveration during reversal and lower repetition index measures for males as compared to females in both phases, and for the 70-30 group as compared to the 90-30 group in the discrimination phase. Plotted is the perseveration index (A), overall repetition index (B), repetition index for the better option (C), and repetition index for the worse option (D) in the discrimination phase. Though we find no significant differences in perseveration index, the 70-30 group shows a significantly lower *RI* and *RI_B_*, and marginally significantly lower *RI_W_*. Additionally, males have significantly lower values for all three repetition index measures. **(E-H)** Same as A-D, but for reversal. We observe a significantly lower perseveration index and a marginally significant lower *RI*, *RI_B_*, and *RI_W_* in males than females. Bars indicate ± *SEM* ^#^p=0.06, *p ≤0.05

**Table 12.** P(stay), p(stay|better), and p(stay|worse) during early discrimination and reversal learning

| Early Discrimination and Reversal Learning |  |  |  |  |  |  |  |
| --- | --- | --- | --- | --- | --- | --- | --- |
| $\gamma \sim [1 + \text{phase} * \text{group} * \text{sex} + (1\vert \text{rats})]$ | | | | | | | |
| $\gamma = P(\text{Stay})$ | | | | | | | |
| Coefficients | $\beta$ | SE | tStat | DF | p | CI<sub>L</sub> | CI<sub>U</sub> |
| intercept | 0.5420 | 0.0054 | 100.37 | 42 | <.0001 | 0.5311 | 0.5529 |
| phase | -0.0172 | 0.0174 | -0.9867 | 42 | 0.3295 | -0.0523 | 0.0179 |
| group | -0.0335 | 0.0234 | -1.4321 | 42 | 0.1595 | -0.0806 | 0.0137 |
| sex | -0.0727 | 0.0426 | -1.7066 | 42 | 0.0953 | -0.1587 | 0.0133 |
| phase:group | 0.0738 | 0.0405 | 1.8211 | 42 | 0.0757 | -0.0080 | 0.1556 |
| phase:sex | 0.0110 | 0.0469 | 0.2351 | 42 | 0.8152 | -0.0836 | 0.1057 |
| group:sex | 0.0434 | 0.0505 | 0.8590 | 42 | 0.3952 | -0.0585 | 0.1453 |
| phase:group:sex | -0.0437 | 0.0623 | -0.7008 | 42 | 0.4873 | -0.1695 | 0.0821 |
| $\gamma = P(\text{Stay}\vert \text{better})$ | | | | | | | |
| Coefficients | $\beta$ | SE | tStat | DF | p | CI<sub>L</sub> | CI<sub>U</sub> |
| intercept | 0.6346 | 0.0143 | 44.351 | 42 | <.0001 | 0.6058 | 0.6635 |
| phase | -0.2521 | 0.0472 | -5.3433 | 42 | <.0001 | -0.3473 | -0.1569 |
| group | -0.0766 | 0.0397 | -1.9307 | 42 | 0.0603 | -0.1566 | 0.0035 |
| sex | -0.1078 | 0.0637 | -1.6919 | 42 | 0.0981 | -0.2363 | 0.0208 |
| phase:group | 0.0172 | 0.0791 | 0.2172 | 42 | 0.8291 | -0.1424 | 0.1768 |
| phase:sex | 0.1202 | 0.0834 | 1.4414 | 42 | 0.1569 | -0.0481 | 0.2886 |
| group:sex | 0.0493 | 0.0784 | 0.6281 | 42 | 0.5333 | -0.1090 | 0.2075 |
| phase:group:sex | 0.0086 | 0.1088 | 0.0787 | 42 | 0.9376 | -0.2109 | 0.2281 |
| $\gamma = P(\text{Stay}\vert \text{worse})$ | | | | | | | |
| Coefficients | $\beta$ | SE | tStat | DF | p | CI<sub>L</sub> | CI<sub>U</sub> |
| intercept | 0.3761 | 0.0219 | 17.134 | 42 | <.0001 | 0.3318 | 0.4204 |
| phase | 0.2285 | 0.0434 | 5.2692 | 42 | <.0001 | 0.1410 | 0.3160 |
| group | 0.0295 | 0.0333 | 0.8868 | 42 | 0.3802 | -0.0377 | 0.0968 |
| sex | -0.0157 | 0.0268 | -0.5869 | 42 | 0.5604 | -0.0697 | 0.0383 |
| phase:group | 0.0267 | 0.0637 | 0.4200 | 42 | 0.6766 | -0.1018 | 0.1553 |
| phase:sex | -0.0718 | 0.0497 | -1.4435 | 42 | 0.1563 | -0.1721 | 0.0286 |
| group:sex | 0.0501 | 0.0437 | 1.1464 | 42 | 0.2581 | -0.0381 | 0.1383 |
| phase:group:sex | -0.0548 | 0.0754 | -0.7274 | 42 | 0.4710 | -0.2069 | 0.0973 |

**Table 13.**
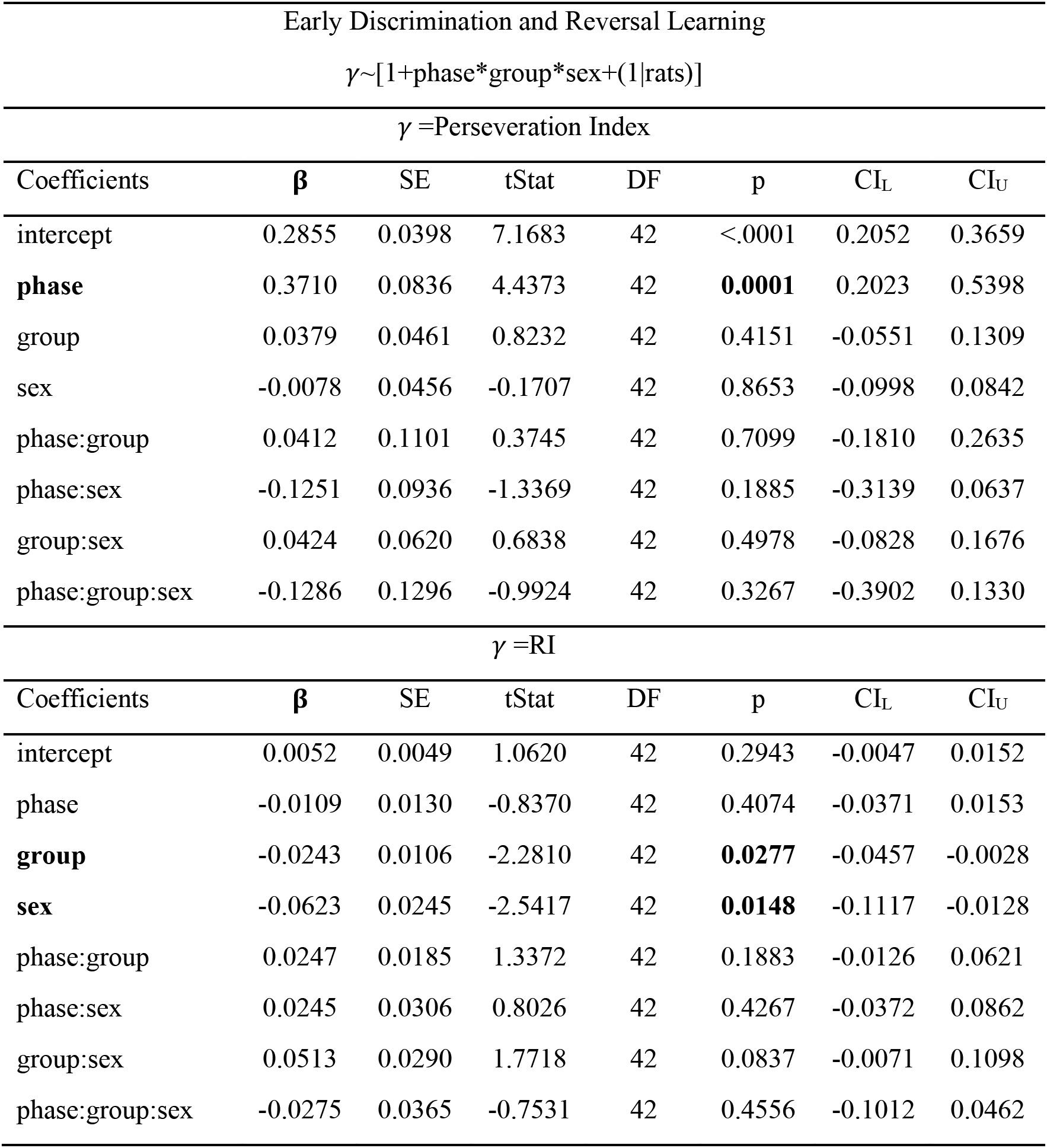
Perseveration index and repetition index (RI) during early discrimination and reversal learning

| Early Discrimination and Reversal Learning |  |  |  |  |  |  |  |
| --- | --- | --- | --- | --- | --- | --- | --- |
| $\gamma \sim [1 + \text{phase} * \text{group} * \text{sex} + (1 \vert \text{rats})]$ | | | | | | | |
| $\gamma = \text{Perseveration Index}$ | | | | | | | |
| Coefficients | $\beta$ | SE | tStat | DF | p | CI<sub>L</sub> | CI<sub>U</sub> |
| intercept | 0.2855 | 0.0398 | 7.1683 | 42 | <.0001 | 0.2052 | 0.3659 |
| **phase** | 0.3710 | 0.0836 | 4.4373 | 42 | **0.0001** | 0.2023 | 0.5398 |
| group | 0.0379 | 0.0461 | 0.8232 | 42 | 0.4151 | -0.0551 | 0.1309 |
| sex | -0.0078 | 0.0456 | -0.1707 | 42 | 0.8653 | -0.0998 | 0.0842 |
| phase:group | 0.0412 | 0.1101 | 0.3745 | 42 | 0.7099 | -0.1810 | 0.2635 |
| phase:sex | -0.1251 | 0.0936 | -1.3369 | 42 | 0.1885 | -0.3139 | 0.0637 |
| group:sex | 0.0424 | 0.0620 | 0.6838 | 42 | 0.4978 | -0.0828 | 0.1676 |
| phase:group:sex | -0.1286 | 0.1296 | -0.9924 | 42 | 0.3267 | -0.3902 | 0.1330 |
| $\gamma = \text{RI}$ | | | | | | | |
| Coefficients | $\beta$ | SE | tStat | DF | p | CI<sub>L</sub> | CI<sub>U</sub> |
| intercept | 0.0052 | 0.0049 | 1.0620 | 42 | 0.2943 | -0.0047 | 0.0152 |
| phase | -0.0109 | 0.0130 | -0.8370 | 42 | 0.4074 | -0.0371 | 0.0153 |
| **group** | -0.0243 | 0.0106 | -2.2810 | 42 | **0.0277** | -0.0457 | -0.0028 |
| **sex** | -0.0623 | 0.0245 | -2.5417 | 42 | **0.0148** | -0.1117 | -0.0128 |
| phase:group | 0.0247 | 0.0185 | 1.3372 | 42 | 0.1883 | -0.0126 | 0.0621 |
| phase:sex | 0.0245 | 0.0306 | 0.8026 | 42 | 0.4267 | -0.0372 | 0.0862 |
| group:sex | 0.0513 | 0.0290 | 1.7718 | 42 | 0.0837 | -0.0071 | 0.1098 |
| phase:group:sex | -0.0275 | 0.0365 | -0.7531 | 42 | 0.4556 | -0.1012 | 0.0462 |

**Table 14.** Repetition index on the better option (RI_B_) and repetition index on the worse option (RI_W_) during early discrimination and reversal learning

| Early Discrimination and Reversal Learning |  |  |  |  |  |  |  |
| --- | --- | --- | --- | --- | --- | --- | --- |
| $\gamma \sim [1 + \text{phase} * \text{group} * \text{sex} + (1 \vert \text{rats})]$ | | | | | | | |
| $\gamma = RI_B$ | | | | | | | |
| Coefficients | $\beta$ | SE | tStat | DF | p | CI<sub>L</sub> | CI<sub>U</sub> |
| intercept | 0.0024 | 0.0023 | 1.0606 | 42 | 0.2949 | -0.0022 | 0.0070 |
| phase | -0.0054 | 0.0064 | -0.8437 | 42 | 0.4036 | -0.0182 | 0.0075 |
| **group** | -0.0121 | 0.0052 | -2.3252 | 42 | **0.0250** | -0.0225 | -0.0016 |
| **sex** | -0.0308 | 0.0122 | -2.5192 | 42 | **0.0157** | -0.0555 | -0.0061 |
| phase:group | 0.0122 | 0.0092 | 1.3214 | 42 | 0.1935 | -0.0064 | 0.0308 |
| phase:sex | 0.0122 | 0.0152 | 0.8033 | 42 | 0.4263 | -0.0185 | 0.0429 |
| group:sex | 0.0259 | 0.0145 | 1.7912 | 42 | 0.0805 | -0.0033 | 0.0551 |
| phase:group:sex | -0.0139 | 0.0182 | -0.7607 | 42 | 0.4511 | -0.0506 | 0.0229 |
| $\gamma = RI_W$ | | | | | | | |
| Coefficients | $\beta$ | SE | tStat | DF | p | CI<sub>L</sub> | CI<sub>U</sub> |
| intercept | 0.0028 | 0.0027 | 1.0606 | 42 | 0.2949 | -0.0025 | 0.0082 |
| phase | -0.0055 | 0.0066 | -0.8296 | 42 | 0.4115 | -0.0189 | 0.0079 |
| **group** | -0.0122 | 0.0055 | -2.2351 | 42 | **0.0308** | -0.0232 | -0.0012 |
| **sex** | -0.0314 | 0.0123 | -2.5632 | 42 | **0.0140** | -0.0562 | -0.0067 |
| phase:group | 0.0126 | 0.0093 | 1.3515 | 42 | 0.1838 | -0.0062 | 0.0313 |
| phase:sex | 0.0123 | 0.0154 | 0.8016 | 42 | 0.4273 | -0.0187 | 0.0433 |
| group:sex | 0.0254 | 0.0145 | 1.7514 | 42 | 0.0872 | -0.0039 | 0.0547 |
| phase:group:sex | -0.0136 | 0.0183 | -0.7451 | 42 | 0.4604 | -0.0506 | 0.0233 |

#### Summary

We found more perseveration during early reversal learning, compared to the discrimination phase. This indicates animals continued to perform according to contingencies learned in discrimination. When using the RI measure, we observed less repetition (lower RI, particularly *RI_W_*) in the 70-30 reward probability group and in males, suggesting probability group and sex have an effect on repetitive behavior that are not due to differences in the propensity to be rewarded.

### Comparisons of Estimated Parameters based on Fit of Choice Behavior with RL models

#### Comparisons between Discrimination & Reversal Learning

To gain more insight into learning and decision making, we next estimated model parameters (learning rate, *α* and sensitivity to difference in subjective reward values, *σ*) based on two RL models across groups in each of the two phases. Unlike our analyses of response to reward feedback and repetition measures, we include all sessions in our RL analysis rather than the first seven to capture learning over a longer time period. We found that the single-learning rate model, RL1, fit the choice data significantly better, as shown by the lower BIC value (difference in BIC between RL1 and RL2 = −7.78, pairwise t-test: t(49) = −75.8, p < 0.0001). For this reason, only results for RL1 are presented below.

Comparison of estimated parameters from RL1 revealed that the average learning rate was significantly higher during reversal than discrimination phase (difference in *α* =0.28; two sample t-test: t(48)=-5.48, p<0.0001) and at the same time, sensitivity to subjective reward values was significantly lower (difference in *σ*=-0.77; two sample t-test: t(48)=8.18, p<0.00001), which corresponds to enhanced exploration. Additionally, to better explore the relationship between learning rate and sensitivity to difference in reward values, we calculated the correlation coefficients between parameters across groups. We found a significant negative correlation between learning rate and sensitivity to difference in reward values during reversal (r=-0.76, p=0.0028) but not discrimination learning (r=-0.42, p=0.0762). We also calculated correlation between model parameters using the inverse Hessian at the ML estimate (see Methods). However, we did not find any evidence that the correlations between model parameters of the RL1 model to be significantly different from 0 during discrimination (r=-0.16±0.20, t(24) = −0.82, p = 0.42) or reversal learning (r=0.017±0.14, t(24) = 0.12, p = 0.90). These results suggest that the observed simultaneous increase in learning rates and decrease in sensitivity to subjective reward value in the reversal relative to discrimination phase was not due to our fitting procedure and, instead, happened due to independent mechanisms.

#### Comparisons within Discrimination Learning

Using estimated parameters based on RL1, we found that the 90-30 reward probability group had a learning rate *α* = 0.038 ± 0.017, sensitivity to difference in reward values *σ* = 0.92 ± 0.12 and fit of *BIC* = 2227 ± 600 (**Figure 7A**). The 70-30 reward probability group had a learning rate *α* = 0.032 ± 0.031, sensitivity to difference in reward values *σ* = 0.86 ± 0.12 and fit of *BIC* = 4234 ± 622. We found no significant differences between reward probability groups in terms of *α* (two sample t-test: *t*(23)= 0.74, p=0.47) or *σ* (two sample t-test; *t*(23)= −0.32; p=0.76). *Comparisons within Reversal Learning.* As in the discrimination phase, we again fit choice behavior for each subject in the reversal phase using RL1 (**Figure 7B**). We found that the 90-30 reward probability group had a learning rate *α* = 0.36 ± 0.069, sensitivity to difference in reward values *σ* = 0.082 ± 0.056 and fit of *BIC* = 3167 ± 390. The 70-30 reward probability group had a learning rate *α* = 0.26 ± 0.061, sensitivity to difference in reward values *σ* = 0.13 ± 0.04 and fit of *BIC* = 5262 ± 396. We found no significant difference between the *α* (two sample t-test; *t*(23)= −0.96; p=0.35) or *σ* (two sample t-test; *t*(23) = 0.70; p=0.49) parameters between groups.

**Figure 7.**
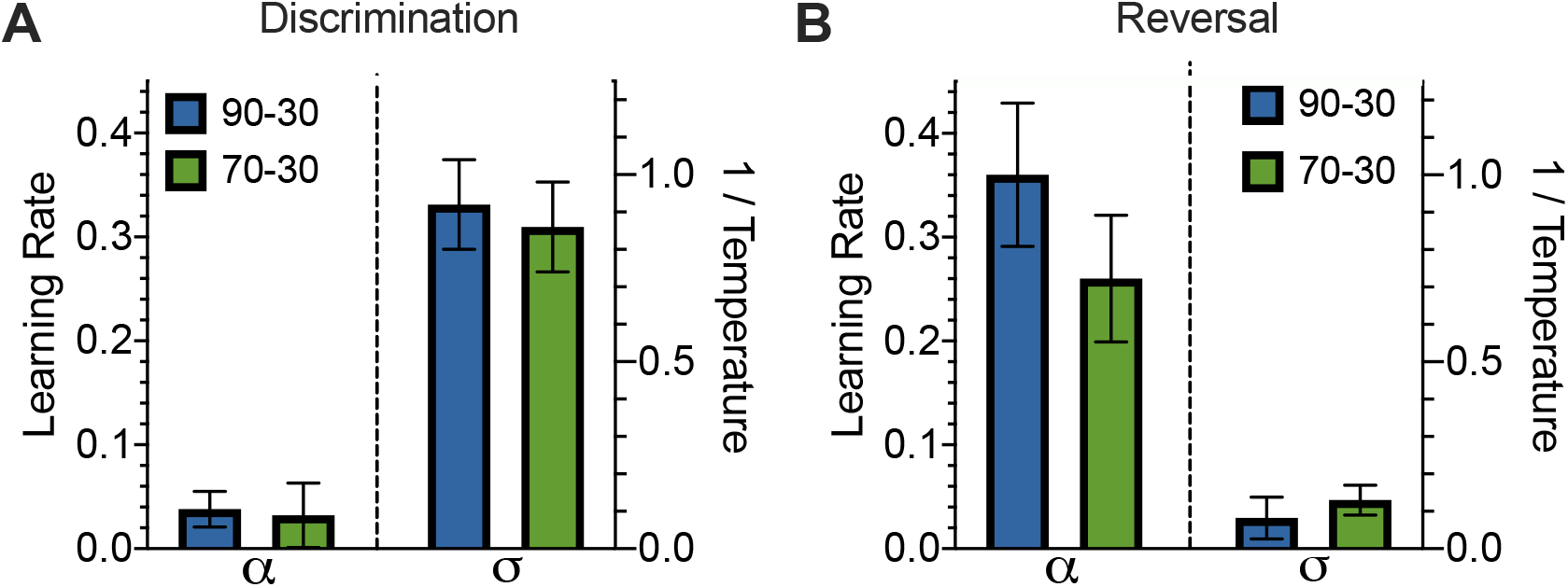
Higher learning rate and lower sensitivity to difference in subjective reward values in reversal compared to discrimination learning. **(A-B)** Learning parameters and sensitivity to difference in reward values for the single-learning rate model during the discrimination (A) and reversal (B) phases. We find no significant difference in parameter values between reward probability groups during discrimination or reversal. However, we do find significantly higher learning rates and significantly lower sensitivity to difference in reward values parameters following reversal.

#### Summary

In analyses across discrimination and reversal phases, we found that sensitivity to difference in reward values was lower during reversal than discrimination, and learning rate(s) were higher (the single learning rate for RL1, and both learning rates for RL2) during reversal relative to the discrimination phase. These results indicate that reversal caused faster learning and more exploration at the same time. However, within each phase of learning, the estimated model parameters were not significantly different between probability groups in either phase. We also found no sex-dependent differences. This suggests the slight increased probability of reward corresponding to the better option in the 90-30 reward probability group, as compared to the 70-30 group, was not large enough to induce significantly different learning or, alternatively, this effect could not be captured by our RL models.

## Discussion

Here, we used a stimulus-based probabilistic discrimination and reversal paradigm with different probabilities of reward (i.e., 90/30, 70/30) to test learning and performance on several measures (i.e., probability of choosing the better option, initiation and choice omissions and latencies, perseveration and repetition measures, and win-stay/lose-shift strategies). We also examined whether fit of choice data using RL models would reveal differences between the two learning phases, and tested for potential differences in learning rates (single, separate) and explore/exploit behavior. We found higher learning rates in the reversal than discrimination phase, which mirror recent reports (described in more detail below). Animals also exhibited decreased sensitivity to the difference in subjective reward values of the two options during the reversal learning phase compared to discrimination, indicative of greater exploration. We discovered consistent patterns across discrimination and reversal learning phases in latencies to choose the better option and latencies to collect reward; the former specific to reversal. Further, we found increased perseveration in early reversal compared to early discrimination learning. Finally, we found that differences in reward collection latencies depended on the richness of the environment (90% vs 70% reward). Notably, the only pronounced sex difference was in longer latencies to choose the better option during reversal learning (females taking longer than males), which is generally consistent with the pattern of effects of a recent study by our group (Aguirre et al., 2020). We elaborate on these findings below.

### Learning rates

We found that the learning rate was higher in reversal than discrimination learning. The general trend of increased learning rates following reversal and decreased sensitivity to the difference in subjective reward values is consistent with existing literature (Costa, Tran, Turchi, & Averbeck, 2015; Massi, Donahue, & Lee, 2018). The increased learning rate suggests a more rapid integration of reward feedback, while the lower sensitivity to difference in reward values indicates rats choose the higher-valued option less consistently, corresponding to greater exploration. These changes may reflect response to increased environmental volatility following the reversal in terms of both faster learning and more exploration. Bathellier, Tee, Hrovat, and Rumpel (2013) accounted for a similar change in learning rate in mice by assuming slower initial learning during discrimination due to weaker initial synaptic weights, and much faster learning during reversal due to the same synapses being activated, but at a state where synaptic weights are stronger than they were in discrimination. Future modeling studies are needed to explain how a single reversal can induce both higher learning rates and decreased sensitivity to subjective values.

### Latencies to collect reward and choose the better option

Probability group differences across both learning phases were found for reward collection latencies, with the 90-30 group exhibiting longer reward collection times than the 70-30 group, suggesting the 90-30 group may experience attenuated motivation by comparison. Latency to collect reward is commonly used as a measure of motivation, whereas stimulus response latency (i.e. latency to choose the better option) is often used as a measure of processing or decision-making speed in the Five-Choice Serial Reaction Time Task (5CSRTT) (Amitai & Markou, 2010; Asinof & Paine, 2014; Bari, Dalley, & Robbins, 2008; Bushnell & Strupp, 2009; Chudasama et al., 2003; Remmelink, Chau, Smit, Verhage, & Loos, 2017; Robbins, 2002; Robinson et al., 2009). One interpretation of this finding is that animals may have been more motivated to confirm whether they had received a reward under higher uncertainty (i.e., when the probability for the better option was lower), and conversely, were more confident in their decision with a higher reward probability associated with choosing the better option. Another related explanation is that animals exert more effort and display more vigor when in leaner and more uncertain reward environments (Amsel, 1967; McNamara, Fawcett, & Houston, 2013), which in our experiments was directly tied to the differences in reward probabilities associated with the better option. Because there was greater reward uncertainty associated with the better option for the 70-30 group, rats may have exerted more effort to retrieve rewards compared to the 90-30 group. Interestingly, although animals generally increased their choice of the better option across days during discrimination learning, this was actually not as much the case for reversal learning where this measure over time was relatively flat. Despite the reward collection latency differences, we found no difference in the number of cached rewards between the probability groups, so these differences are likely not due to satiety *per se*, but instead, an effect of an experience with overall reward rate over time.

A final consideration related to our consistent probability group difference is a recent report (Song & Lee, 2020) providing evidence for differential neural recruitment in environments that require different “resource allocations.” Agents (or in our case, rats) learn over time to assign resources to certain stimuli, and make adjustments as stimuli gain more reward-predictive value. The leaner reward schedule (70-30) may actually promote a more adjustable, flexible resource allocation than the more profitable schedule (90-30). Empirically testing how midbrain dopamine interacts with cortical structures to support flexible resource allocation is an interesting line of future inquiry. To probe this, different reward probability schedules could be compared.

### Perseveration and Win-Stay Strategies

All animals exhibited more perseveration during early reversal learning than early discrimination learning. This finding supports those by Verharen, den Ouden, Adan, and Vanderschuren (2020) in which they simulated data of thousands of probabilistic reversal learning sessions using a Q-learning model consisting of separate learning rates for learning from positive (i.e. rewarded) and negative (i.e. nonrewarded) feedback, a beta parameter, and a stickiness parameter indicative of perseveration. They found that a greater number of reversals occurred when the stickiness parameter value was high (i.e., greater perseveration), but only when both learning rates were also high (Verharen et al., 2020). Interestingly, we found reward probability group differences for perseveration, with the 90-30 group exhibiting greater perseveration than the 70-30 group. This finding implies that more consistent reward feedback (i.e. higher reward probability associated with the better option) in the 90-30 group promoted perseverative responding in early learning. Furthermore, males demonstrated greater exploratory behavior (i.e. lower perseveration) during discrimination learning, in line with previous research showing greater impulsivity in male rats (Lukkes, Thompson, Freund, & Andersen, 2016; Palanza, Gioiosa, & Parmigiani, 2001).

Animals will perseverate on stimulus choice during *early* reversal learning, yet animals eventually alternate and switch according to the new reward contingencies over time. Greater perseveration during early reversal is consistent with the long-standing idea that reversal learning is a measure of inhibitory control, such that the inability to disengage from previously rewarding behavior after a change in contingency may be reflective of compulsive and even impulsive response tendencies, commonly associated with drug dependence (Izquierdo & Jentsch, 2012). Indeed, there is evidence that inflexible responding in reversal learning may be genetically related to impulsivity (Crews & Boettiger, 2009; Fineberg et al., 2010; Franken, van Strien, Nijs, & Muris, 2008; Groman, James, & Jentsch, 2009; Groman & Jentsch, 2013; Jentsch et al., 2014). However, greater perseveration can also be explained by slower updating of model-free learning in which animals are not benefiting as much from trial-by-trial feedback, or (just as likely) failing to detect task transitions, especially in early reversal (Izquierdo et al., 2017). It is particularly interesting that we show here *stimulus* perseveration, aside from spatial- or response-based perseveration, which we control for here by pseudorandomly presenting the stimuli on the left- vs. right-sides of the screen.

A related observation we report here is that animals more often used reward-dependent choice strategies (i.e., win-stay and win-stay|better) in the early discrimination phase compared to the reversal phase, and adopted an opposing pattern after reversal (i.e., win-stay|worse as more prevalent). Importantly, as above, this strategy is stimulus-dependent and not location-dependent, which is more often probed in lever-based (left vs. right) tasks. Indeed, animals exhibited less consistent response to reward feedback during the reversal compared to the discrimination phase, indicative of noisier behavior or equivalently more exploration.

### Sex differences

Females exhibited longer latencies to choose the better option than males, and this was only observed during the reversal learning phase. Measures of decision speed in rodents vary, but can include latencies to nosepoke (i.e., time to make a response before the end of the trial), as well as percentage of correct responses (i.e., accuracy), usually on the 5CSRTT as described above. Our data supports the interpretation that females exhibit greater response demands (and consequently slower performance) than males in the reversal phase. This adds to a growing literature on sex differences in reversal learning (Aarde et al., 2020; Bissonette et al., 2012; Branch et al., 2020; LaClair & Lacreuse, 2016).

### Conclusion

In conclusion, we outline ways in which discrimination and reversal learning are behaviorally more unique than they are similar: sex differences in latencies to choose the better option, lesser Win-Stay strategies employed, and more perseveration in early reversal learning. Additionally, fitting choice behavior with reinforcement learning models, we found a lower sensitivity to the difference in subjective reward values (greater exploration) and higher learning rate for the reversal phase compared to original discrimination learning phase. Interestingly, a consistent reward probability group difference emerged with a richer environment associated with longer reward collection latencies than a leaner environment. We also replicate previous reports on sex differences in reversal learning. Together, our results suggest that metrics of decision speed and motivation should be used as more than auxiliary measures to study reversal learning. Future studies should probe the neural correlates of these fine-grained behavioral measures, as these have been under-utilized and may reveal marked dissociations in how the circuitry is recruited in each phase.

## Supplemental Figure

**Figure S1.**
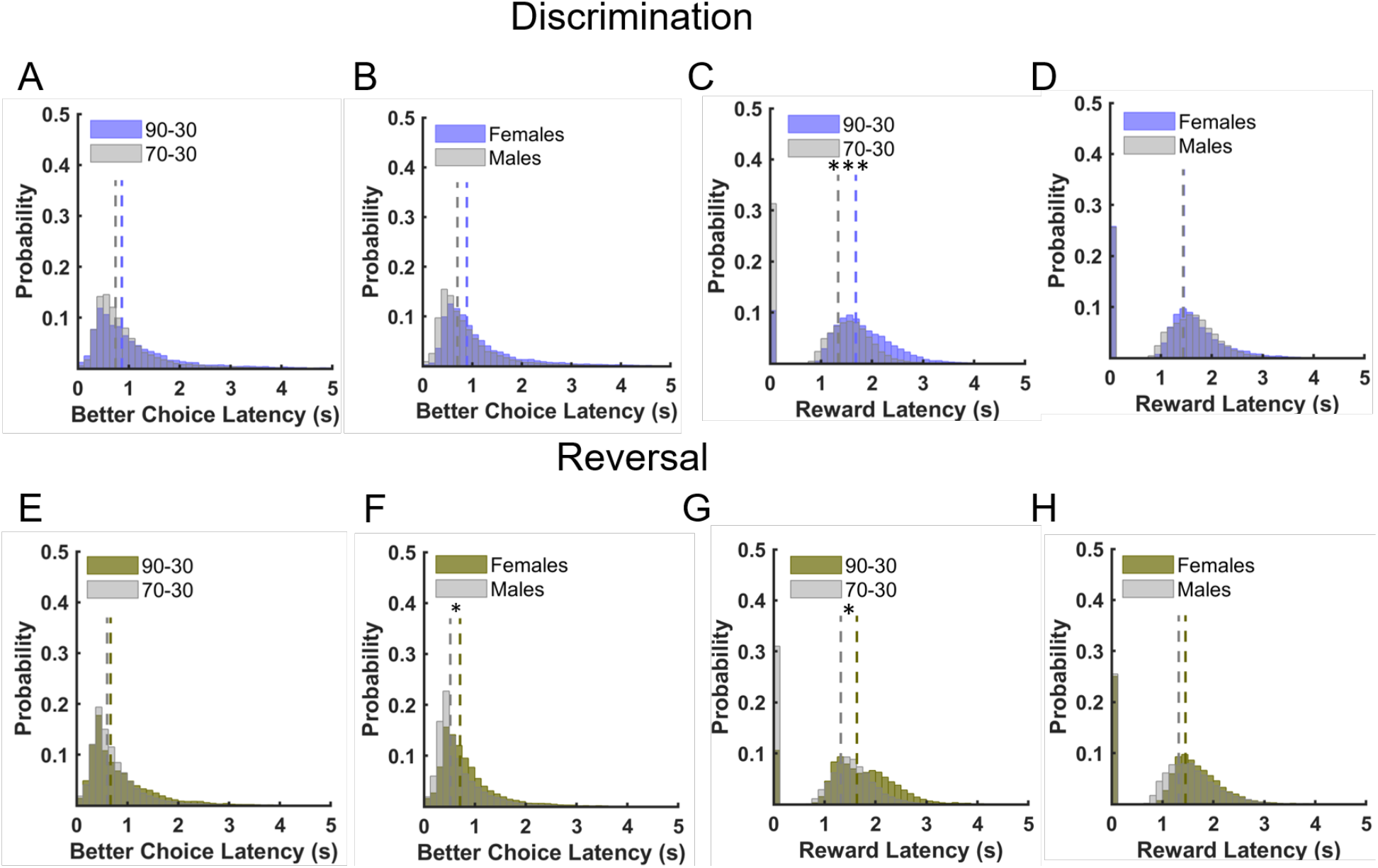
90-30 probability group was consistently slower to collect reward than 70-30 probability group. **(A-B)** There were no group or sex differences in better choice latencies for discrimination learning. The 90-30 group exhibited longer reward collection latencies than the 70-30 group in discrimination learning. **(D)** No significant sex differences in reward collection latencies in discrimination learning. **(E)** No significant group differences in better choice latencies in reversal learning. **(F)** Females exhibited longer latencies for the better option in reversal learning, with or without controlling for discrimination sessions to criterion. **(G)** The 90-30 group exhibited longer reward collection latencies than the 70-30 group in reversal learning, with or without controlling for discrimination sessions to criterion. (**H**) No significant sex differences in reward collection latencies in reversal learning. Dashed lines in histograms of latencies represent group medians. *p ≤0.05, ***p≤0.001

## Acknowledgements

We thank Alexandra Stolyarova for programming the probability schedules in the Lafayette Instrument chambers and for statistical consultation on General Linear Models.

